# Anisotropic Cellular Mechanoresponse for Radial Size Maintenance of Developing Epithelial Tubes

**DOI:** 10.1101/172916

**Authors:** Tsuyoshi Hirashima, Taiji Adachi

## Abstract

Cellular behaviors responding to mechanical forces control the size of multicellular tissues as demonstrated in isotropic size maintenance of developing tissues. However, how mechanoresponse systems work to maintain anisotropic tissue size including tube radial size remains unknown. Here we reveal the system underlying radial size maintenance of the murine epididymal tubule by combining quantitative imaging, mathematical modeling, and mechanical perturbations. We found that an oriented cell intercalation making the tubule radial size smaller counteracts a cell tension reduction due to neighbor cell division along the tubule circumferential axis. Moreover, we demonstrated that the tubule cells enhance actomyosin constriction driving the cell intercalation in response to mechanical forces anisotropically applied on the cells. Our results suggest that epididymal tubule cells have endogenous systems for responding as active cell movement to mechanical forces exclusively along the circumferential axis, and the anisotropic cellular mechanoresponse spontaneously controls the tubule radial size.

## Introduction

Cell-cell communication via cellular behaviors is necessary to regulate homeostasis including size maintenance in development. The underlying principle for this communication is a cellular response in reaction to mechanical forces generated within the tissues (Guillot and Lecuit, 2013; Mammoto et al., 2013). In tissue volume maintenance, for example, cell proliferation is counterbalanced by cell death, and these coordinated cellular behaviors are invoked in response to cell-crowding pressure generated by cell proliferation as a mechanoresponse (Eisenhoffer et al., 2012; Gudipaty et al., 2017; Hufnagel et al., 2007; Shraiman, 2005). On the other hand, in tissue length maintenance along an axis, such as radial size maintenance of tubes during elongation (Andrew and Ewald, 2010; Hirashima, 2014; Karner et al., 2009), anisotropic mechanoresponse systems in tissues might organize cooperative cellular behaviors to distribute a net increase in the tissue volume not along the axis but in the other axes.

The length maintenance in developing tissues can be considered as another aspect of anisotropic tissue growth giving rise to a variety of tissue shape (Thompson, 1942). Recent studies have revealed so far that mainly two factors at cellular level underpin the anisotropic tissue growth. One is cell division orientation, determined by the position of the mitotic spindle. It has been shown that the shape of growing tissues largely obeys the orientation of cell division, and also increasing evidence implicates the misorientation of cell division leads to the abnormal morphology in tissues as demonstrated in various organs and species (Fischer et al., 2006; Morin and Bellaïche, 2011; Tang et al., 2011). The other one is cell intercalation, active cellular movement with changing the position of cell junctions in neighboring cells (Guillot and Lecuit, 2013; Walck-Shannon and Hardin, 2014). It has been clearly demonstrated that the cell intercalation is driven by polarized myosin II localization at cell-cell junctions and cumulative effect of the local tissue dynamics results in anisotropic large deformation at the tissue scale (Bertet et al., 2004; Nishimura et al., 2012; Zallen and Blankenship, 2008). These cellular events generate mechanical forces and thereby impact on surrounding cells in multicellular packing tissues, suggesting a mechanoresponse system interconnecting constituent cells would be formed in developing tissues. However, it remains unclear how the system works to be responsible for the tissue length maintenance in the dynamics.

Here we utilized epididymal tubules in murine embryo as an experimental model system to reveal anisotropic cellular responses to mechanical forces. The epididymal tubule is a single mono-layered epithelial tube that shows dynamic morphological change including bending and folding during the development from embryonic day (E) 15.5 (Hirashima, 2014; Hirashima, 2016; Joseph et al., 2009). Interestingly, the radial size of developing epididymal tubule is almost constant even though the tissue volume increases due to incessant cell proliferation in the tubules (Hirashima, 2014; Hirashima and Adachi, 2015; Tomaszewski et al., 2007). To reveal the mechanism to explain this phenomenon, Xu et al. have shown that the oriented cell intercalation is involved in the tubule elongation with maintaining the radial size (Xu et al., 2016). If the oriented cell intercalation have sustained in the epididymal tubule, the radial size would become smaller along the time during the development, inconsistent with the observation. Therefore, this study motivates us to unravel unknown regulations for the radial size maintenance of epididymal tubules. In this paper, we employed interdisciplinary approach combining quantitative imaging and mathematical approaches, and found that the epididymal cells counteract mechanical forces exclusively along the circumferential axis of tubule. We finally propose the anisotropic mechanoresponse system at supra-cellular scales that operates to maintain the tubule radial size at whole tissue scale.

## Results

### Non-oriented cell division produces mechanical forces to expand the tubule

Since the cell division orientation has been known as a key determinant for the morphogenesis of developing tubes (Fischer et al., 2006; Tang et al., 2011), we examined the impact of cell division orientation in the tubule at E15.5 and E16.5. We first performed immunostaining for phospho-histon H3 (pHH3) and γ-tubulin each as a marker of mitotic cells and microtubule organizing centers (MTOCs), and marked spindle orientation in the mitotic cells within the tubule by extracting the axis connecting between the two MTOCs in each pHH3-positive cell without destructing the tissue structure (Fig. 1A-A´´). Then, we manually measured three angles *α*_1_, *α*_2_, and *α*_3_ (Fig. 1B) to define a local coordinate system *O*’ (Fig. 1C), origin of which corresponds to the center position of mitotic cell (white arrows in Fig. 1B). In the local sphere coordinates, we obtained statistical distributions for two angles of the cell division orientation: *φ* and *θ* (Fig. 1C; Materials and Methods (II-i)). The distribution of *φ* shows that most of the mitotic cells divide parallel to the epithelial layer (The Rayleigh test, *P*<0.001 at E15.5 and E16.5); however, the distribution of *φ* shows that the cell division orientation is not biased towards the longitudinal or circumferential axis (The Rayleigh test, *P*>0.05 at E15.5 and E16.5) (Fig. 1D). Furthermore, the distance between MTOCs shows no correlations with the angle *θ*(*r*=−0.07 at E15.5, *r*=−0.02 at E16.5 where *r* is the Pearson's correlation coefficient) (Fig. 1E), rejecting that the spindle orientation might turn to a specific orientation in the later stage of mitosis.

We then investigated how the cell division orientation affects the tubule morphology using numerical simulations of a computational model (Mov. 1). We here employed the vertex dynamics model, a type of cell-oriented model, to represent multi-cellular dynamics in epithelial tubes (Fletcher et al., 2014; Nagai and Honda, 2001; Rauzi et al., 2008). See the Materials and Methods (III-i) - (III-iii) for more details of the mathematical model and simulation settings (Fig. S1). In the simulation, virtual epithelial tubes grew along the division orientation, indicating that cell division along the circumferential axis is attributed to continuous radial growth (Fig. 1F and F´). In addition, we found that the cell division orientation affects mechanical stress in each cell of the virtual tubes (Fig. 1F). To evaluate the mechanical stress in the tube cells, we introduced a scalar quantity, i.e., cell tension anisotropy, which becomes larger when the cell tension along the circumferential axis becomes weaker than that along the longitudinal axis. See Eq. (iv-2) in the Materials and Methods (III-iv) for the definition of cell tension anisotropy. As shown in Figure 1F-F´´, the cell tension anisotropy in the tubes depends on the cell division orientation and corresponds with the growth direction of the tubes. The simulation analysis indicates that the circumferential cell division entails the increase of cell tension anisotropy locally in the tissue, resulting in the tube expansion. This implies that cells should actively counteract the changes in cell tension anisotropy for the maintenance of tubule radial size.

**Figure 1.**
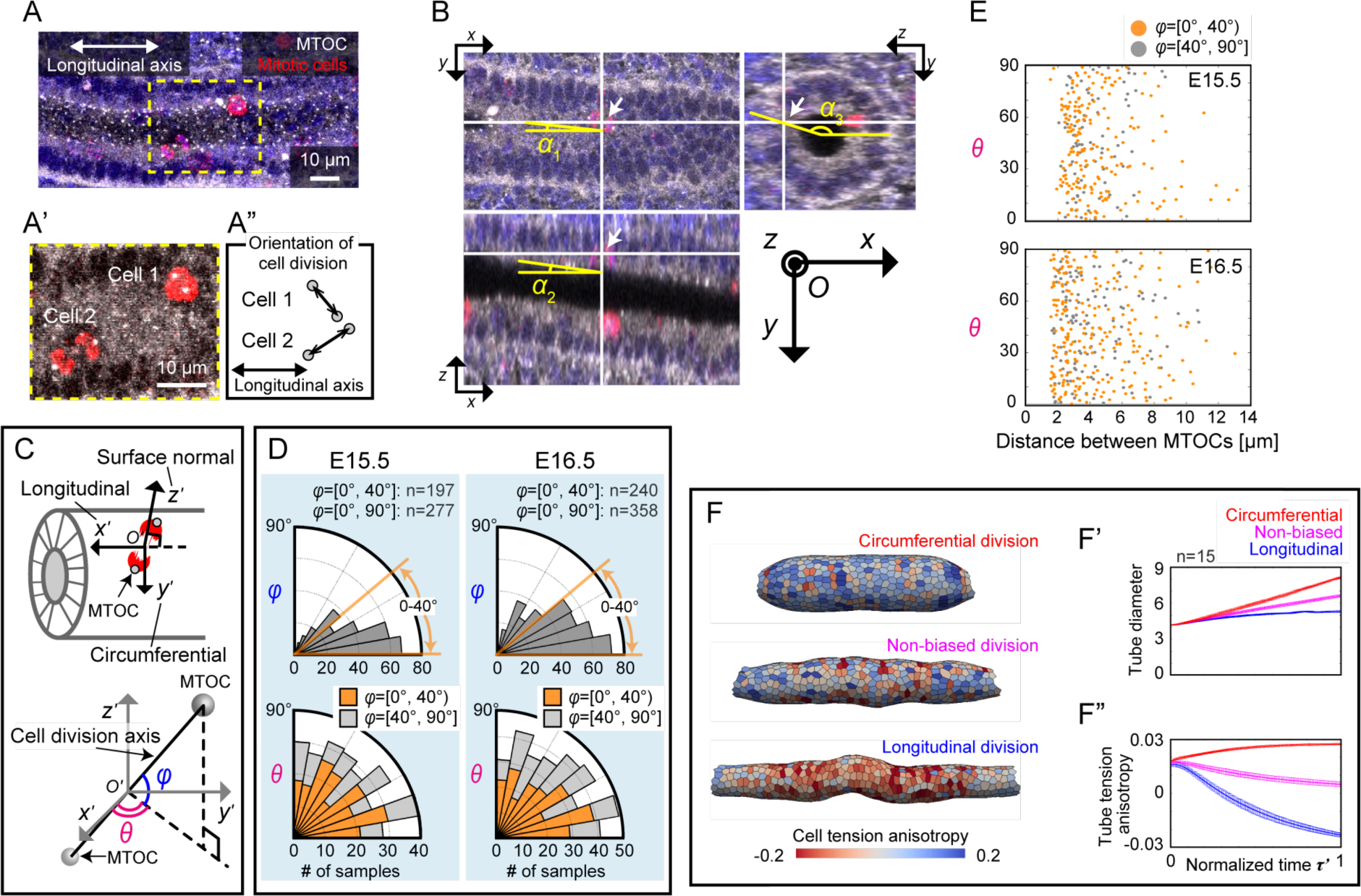
Cell division orientation determines the mechanical stress and tubule morphology. (A) Maximum intensity projection (MIP) of immunostained images for the pHH3 (mitotic cells, red) and γ-tubulin (MTOCs, white). (A´ and A´´) Magnified view of the dotted square in (A), displaying the two cells divide in different orientations. Scale bars: 10 μm. (B) Measurement of cell division orientation in the tubule. Arrows indicate the center position of the mitotic cell. Using the three angles *α_1_,α_2_* and *α_3_*, the observation coordinate *O* is transformed into the local coordinate *O´* in Fig. 1C. (C) Illustration of the local coordinates. (D) The angle distributions of the cell division orientation. Colors in the *θ* distribution represent samples, of which *φ* ranges from 0-40 degrees (orange) and from 40-90 degrees (gray). (E) Scatter plot for the distance between MTOCs and the angle *θ*. The sample number is the same as in (D). (F-F´´) Model simulations in different cell division orientations. (F) The vertex dynamics model simulation of proliferating tubes. The color represents the cell tension anisotropy. (F´ and F´´) Time course of morphological and mechanical quantities along the normalized simulation time *τ*′. See the Materials and Methods (III-iv) and (III-v) for those quantities and Materials and Methods (III-ii) for the definition of *τ*′. *n*=15.

### Polarized actomyosin constriction generated in the tubule cells suppresses the tubule expansion

To study how the cells react against the circumferential cell division, we focused on phosphorylated myosin regulatory light chain (pMRLC), a marker of active tension generation via actomyosin constriction. We first examined the localization of pMRLC in the epididymis by whole-tissue co-immunostaining for the pMRLC and an apical tight junction marker Zonula occludens-1 (ZO-1). The immunostainings revealed that the pMRLC localizes only at apical cellular junctions of the epithelial tubules at E15.5 and E16.5 (Fig. 2A). Furthermore, the pMRLC localization is polarized along the circumferential axis of the tubules (Fig. 2A).

**Figure 2.**
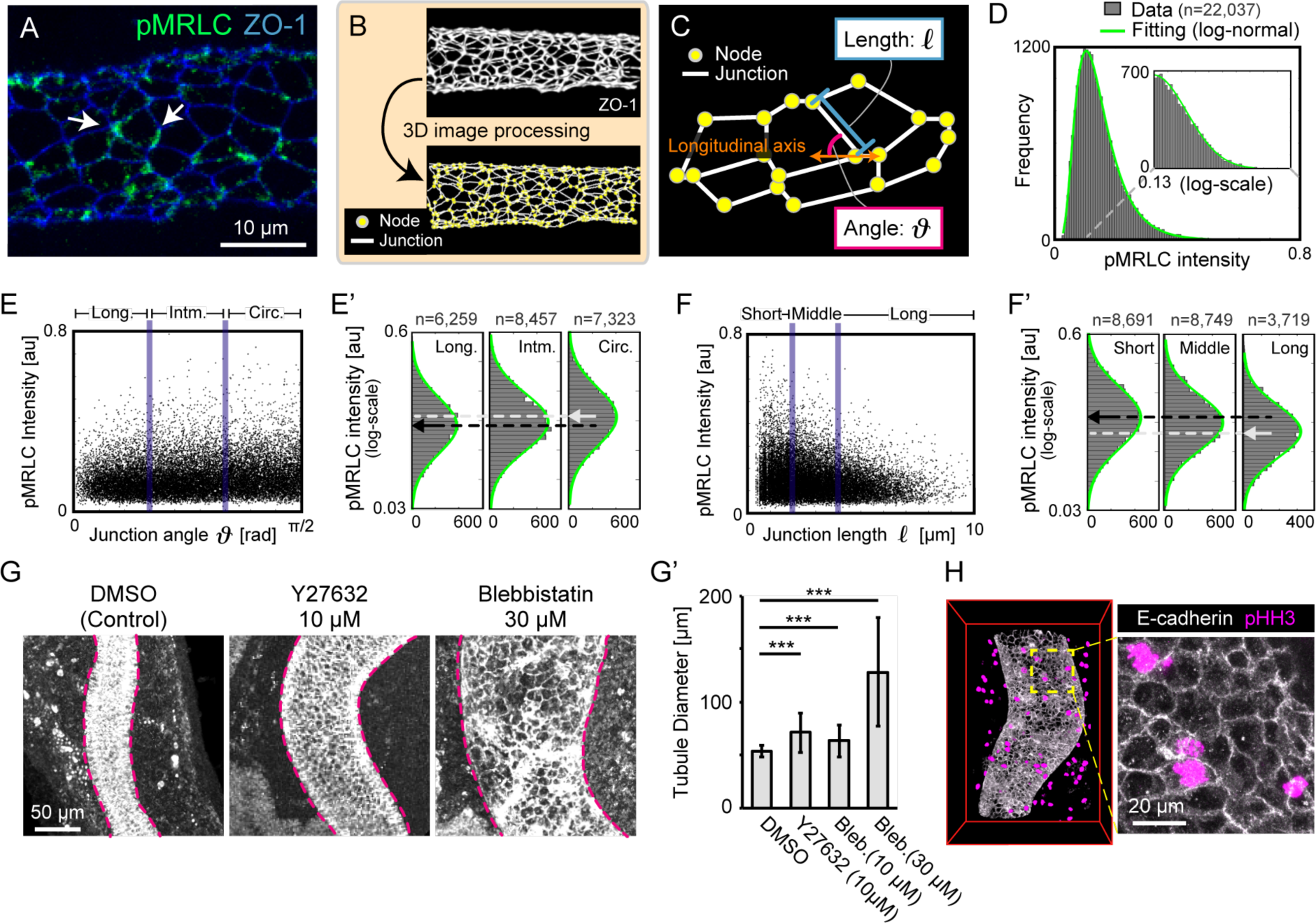
Polarized apical junction constriction is required for the maintenance of tubule radial size. (A) MIP of co-immunostained images of pMRLC (green) and ZO-1 (blue). Arrows indicate pMRLC localization at circumferential apical junctions. Scale bar: 10 μm. (B) Digital processing from immunofluorescence images for the ZO-1 detects nodes and junctions of apical surfaces. (C) Illustration of the junction length *ℓ* and junction angle *ϑ* that measures the angle from the longitudinal axis of tubules. (D) Histogram of pMRLC intensity (gray) fitted with a log-normal distribution (green). The small window displays the tail of distribution. *n*=22,037. (E-F´) Relationship between the pMRLC intensity and the junction angle/length. The log-scale histograms are shown in three divided groups. Black arrows represent the mean intensity in the longitudinal/short group and white arrows represent that in the circumferential/long group. (G and G´) Morphological change of tubules in the inhibitor treatment. (G) MIP of immunofluorescence images for the E-cadherin of head region of epididymis. Scale bar: 50 μm. Magenta dotted lines represent the basal line of tubules in the MIP images. (G´) The tubule diameter for the treatments. *n*=320. (H) 3D images of the co-immunostaining for E-cadherin (white) and pHH3 (magenta) in 30 μM Blebbistatin treatment. Scale bar: 20 μm.

For quantification of the pMRLC spatial distribution at sub-cellular resolution, we measured the pMRLC mean intensity at junctions, junction angle *ϑ*, and junction length *ℓ* through automatic extraction for each apical cell junction by digital image processing (Fig. 2B and C; Materials and Methods (II-ii)). The distribution of pMRLC intensity was well fitted to the heavy-tails of log-normal distribution (Fig. 2D), indicating that enriched pMRLC localizations can be observed in an unignorable number of apical junctions instead of random fluctuations. This statistical distribution suggests that the myosin would be able to be activated by biological events that occur intermittently with low frequency rather by constant regulation. To evaluate the relationship between the junction angle and the pMRLC intensity, we categorized the junction angle into three groups; longitudinal (long.): 0 ≤*ϑ* ≤*π*/6, intermediate (intm.): *π*/6 <*ϑ* <*π*/3, and circumferential (circ.): *π*/3 ≤*ϑ* ≤*π*/2 (Fig. 2E). As shown in Figure 2E´, the pMRLC tends to localize more on junctions that orient more in the circumferential axis (One-way ANOVA, *P*<0.001). The variation in the position of pMRLC distribution between the junction angles does not show huge difference but reflects that the myosin activation depends on rare events rather than regulations driving large deformation of epithelial tissues such as fly wing or neural tubes. As well as the junction angle, we also categorized the junction length into three groups: short: 0 ≤*ℓ* ≤2 μm, middle: 2 <*ℓ* <4 μm, long: 4 ≤*ℓ* ≤10 μm (Fig. 2F), and found that the pMRLC tends to localize more on shorter junctions (One-way ANOVA, *P*<0.001) (Fig. 2F´). These quantifications suggest that the polarized actomyosin-mediated constriction would shrink the circumferential apical junctions.

We then examined the mechanical roles of actomyosin constriction for the tubule morphology in the inhibitor assay. We applied Blebbistatin/Y-27632 for inhibiting actomyosin constriction by non-muscle myosin II (Kovács et al., 2004; Watanabe et al., 2007) to the epididymides in explant cultures, and observed the tubule morphology 1 day after the inhibitor treatment. The assay revealed that inhibiting actomyosin-mediated apical constriction resulted in the tubule expansion (One-way ANOVA, *P*<0.001) (Fig. 2G and G´; Materials and Methods (II-iii)). In addition, we found that even when the inhibitors were treated, the mitotic cells remain in the epididymal tubules to the same extent as the control ones (Fig. 2H). Although the inhibitor treatment might impede the cytokinesis, the epididymal tubule cells could enter the M phase in the cell cycle. Hence, the inhibiting actomyosin constriction affects little the change in local tissue volume within 1 day. These results indicate that the active constriction polarized circumferentially at the apical junction of epididymal tubules is required for the maintenance of tubule radial size.

### Cell division should trigger the polarized myosin activation for the maintenance of tubule radial size

It has been clarified that the polarized pMRLC localization drives cell intercalation mediated through junction shrinkage in the developing epididymis (Xu et al., 2016) as well as shown in other contexts (Andrew and Ewald, 2010; Guillot and Lecuit, 2013; Walck-Shannon and Hardin, 2014). Hence, the polarized pMRLC localization on the circumferential junctions can drive the oriented cell intercalation, eventually leading to longitudinal elongation and radial shrinkage of the tubule (Bertet et al., 2004; Honda et al., 2008). In contrast, non-oriented cell proliferation results in the tubule expansion as shown in the earlier section. Therefore, the maintenance of tubule radial size requires a proper balance between myosin activity driving intercalation and cell proliferation frequency.

To evaluate this relationship, we measured the pMRLC intensity along the tubule at the whole organ scale at E15.5 and E16.5 (Materials and Methods (II-iv)), and found that the intensity in the head region of epididymis is larger than that in the tail region (Fig. 3A). This intensity profile is similar to the spatial profile of the mitotic cell number in the epididymal tubules previously reported (Hirashima and Adachi, 2015) (Fig. 3B). Indeed, the pMRLC intensity and the mitotic cell numbers in the tubules show positive correlation (*r*=0.49 at E15.5 and *r*=0.56 at E16.5 where *r* is the Pearson's correlation coefficient) (Fig. 3B and B´; Materials and Methods (II-v)). Western blotting analysis also shows the myosin activity, defined as the ratio of pMRLC to the total MRLC (tMRLC), in the head region is greater than that in the tail region (Fig. 3C and C´). Moreover, cell morphology differences corroborate junctional tension due to the pMRLC in each region (Fig. 3D and S2; Materials and Methods (II-vi)). The apical junction length along the circumferential axis in the head region is shorter than that in the tail region (Kruskal-Wallis test, *P*<0.001) (Fig. 3D´), and the apical surface area in the head region are smaller than that in the tail region (Kruskal-Wallis test, *P*<0.001) (Fig. 3D´´).

**Figure 3.**
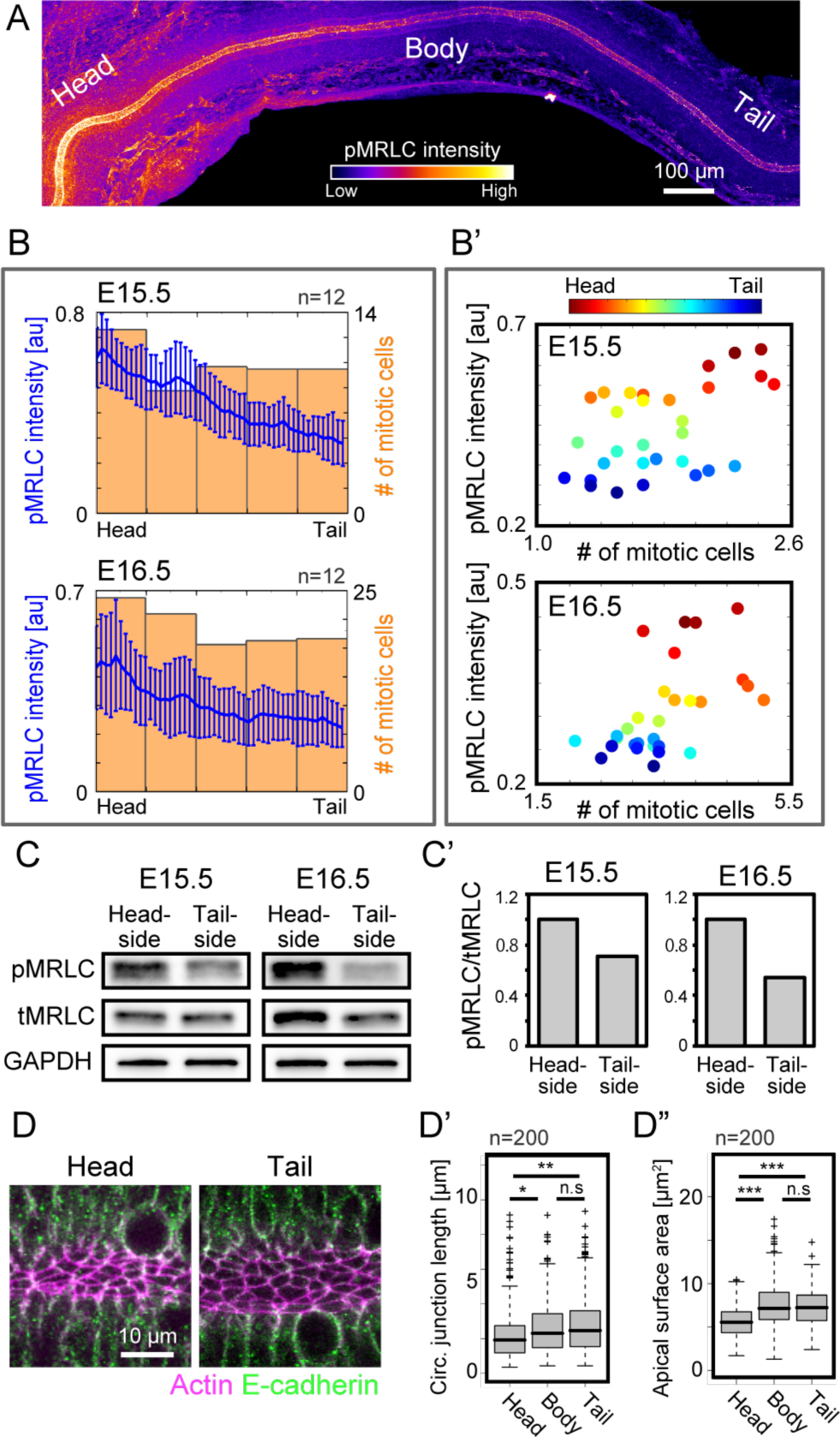
Apical junctional constriction and cell division shows positive correlation. (A) MIP of pMRLC immunostaining at the whole-tissue scale. Scale bar: 100 μm. (B) Spatial profile of pMRLC intensity (blue) and the number of mitotic cells (orange) along the tubule. *n*=12. (B´) The pMRLC intensity versus the mitotic cell number. The color indicates position of data in the tubule. (C and C´) Myosin activity quantification by immunoblotting. (D-D´´) Apical junction morphology visualized by staining for the F-actin and E-cadherin. *n*=200. Scale bar: 10 μm.

Although there are positive correlations between the myosin activity and mitotic cell number in the tubule, the cell proliferation is of no significant dependence on the myosin activity because the inhibition of myosin activity does not affect the mitotic cell number. Thus, it is reasonable to consider that the myosin activation is triggered by the cell proliferation to achieve the balance for the radial size maintenance.

### Cell division yields mechanical stimulations to the neighboring cells

To explore how the cell proliferation triggers the myosin activation, we examined the mechanical effects of cell proliferation in the tubules using the vertex dynamics model. It has been shown that the cell division can produce mechanical perturbations in multi-cellular tissues (Stewart et al., 2011); hence, we anticipated that the cell division apply the pushing forces to the neighbors of divided cells and this stimulus would trigger the polarized myosin activation in the pushed cells. We investigated the change of the cell tension anisotropy against the cell division in the extensive numerical analysis.

We first examined how the cell division affects the cell tension anisotropy in the neighbors of divided cells. To do so, we noticed each spatial and temporal factors: relative angle of target cells to the cell division axis (Fig. 4A) and normalized elapsed time from the cell division 
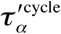
, ranged from 0 to 1 (see the Materials and Methods (III-iii) for the definition). We then evaluated the distribution of cell tension anisotropy in the two cases of cell division orientations, i.e., longitudinal ( 0 ≤*θ* ≤*π*/6 where *θ* is defined in Fig. 1C) and circumferential (*π*/3 ≤*θ* ≤*π*/2) (Fig. 4B). The results shown in Figure 4B´ indicate that the smaller the relative angle, the larger the cell tension anisotropy in the neighbors of divided cells can be altered (upper rows in Fig. 4B´). Note that small relative angle means that the measured cells are located to the circumferential direction of divided cells for the circumferential cell divisions while ones are located to the longitudinal direction of divided cells for the longitudinal cell divisions. This result is intuitive because the divided cells push along the division axis to the neighboring cells, and hence, tension in the neighboring cells along the division axis can be attenuated. In addition, the histograms clearly show that the tension attenuation due to the cell division is the most significant soon after the cell division (the leftmost column in Fig. 4B´) and becomes smaller along with the elapsed time from the cell division 
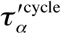
.

**Figure 4.**
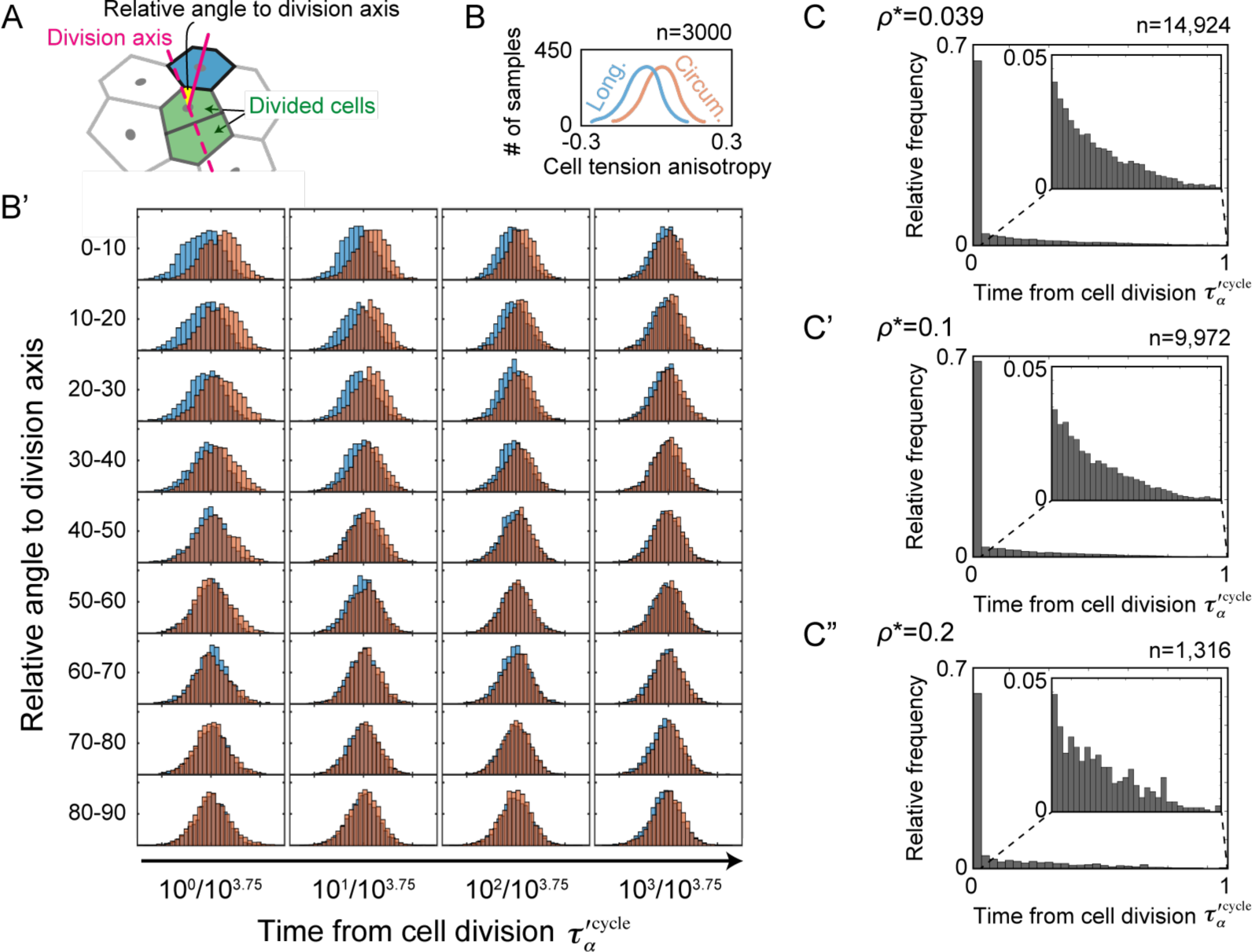
Computational analysis of cell stress influenced by cell division in tubes. (A) The schematics for glossary. The cell tension anisotropy was calculated in cells (e.g., colored in blue) neighboring to the divided cells (green). (B and B´) Histograms of cell tension anisotropy against the cell divisions. (B) The explanatory illustration for the histograms. Red indicates the case of circumferential cell division and blue indicates the case of longitudinal cell division. (B´) Histograms of the cell tension anisotropy in various cases of the relative angle to the cell division axis (vertical) and in those of the elapsed time of neighbors' cell division 
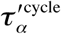
 (horizontal). (C-C´´) Histograms of the elapsed time after the cell division 
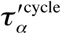
 when the cell tension anisotropy is over various thresholds ( *ρ*\* =0.039, 0.1, and 0.2) in neighboring cells on the circumferential axis. The tail of histograms is magnified in the small windows.

We then examined the time-dependency of change in the cell tension anisotropy from the cell division. In so doing, we monitored cell tension anisotropy *ρ_α_* in all cells, and marked cells as the target cells when *ρ_α_* was over various thresholds *ρ\**. Then, we counted the elapsed time from the cell division 
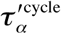
 in the neighboring cells located on the circumferential axis from the target cells, since we have focused on the circumferential cell division that conduces to the tubule expansion. We here set the thresholds as *ρ\** = 0.039, 1, and 0.2 ( *ρ\** = 0.039 is the mean of cell tension anisotropy in the upper-left case of Fig. 4B´) (Fig. 4C-C´´). Our numerical investigations revealed that the cell tension anisotropy in most of the cells (~70%) was altered immediately following to the cell division, consistent with the results of the previous analysis (Fig. 4B´). In addition, we found that the cell tension anisotropy could become over the given thresholds at timing elapsed enough from the cell division (small windows in Fig. 4C-C´´). This occurs because anisotropic deformation of individual cells can be induced by the mechanical stress accumulated in the local regions owing to non-adjacent cell proliferation instead of adjacent cell proliferation.

These analyses provided the two suggestions. First, most of the cells that receive significant mechanical impact from the neighboring divided cells can be found soon after the cell division. Second, cell tension can be varied independent to the cell division, rationalizing the introduction of mechano-responsive regime into our model argued in the later section.

### Mechanical stimulations provided by neighbor cell divisions trigger the onset of cell intercalation

To directly observe cellular responses to the mechanical forces generated by the cell divisions, we performed live imaging analysis for epididymal tubule cells dissected at E15.5 using ex vivo organ culture systems. For visualization of the tubule cell membrane, we crossed the R26R-Lyn-Venus line (Abe et al., 2011) and the Pax2-Cre line (Ohyama and Groves, 2004) to create a conditional fluorescence reporter line. Because the epididymal tubules were located in deep region of the epididymis, ~100 μm away from the capsule of epididymides (Fig. 5A), confocal microscopy was not appropriate for this case; hence, we employed an incubator-integrated multiphoton excitation microscope system for stable deep tissue live imaging in explant cultures (Fig. 5B).

**Figure 5.**
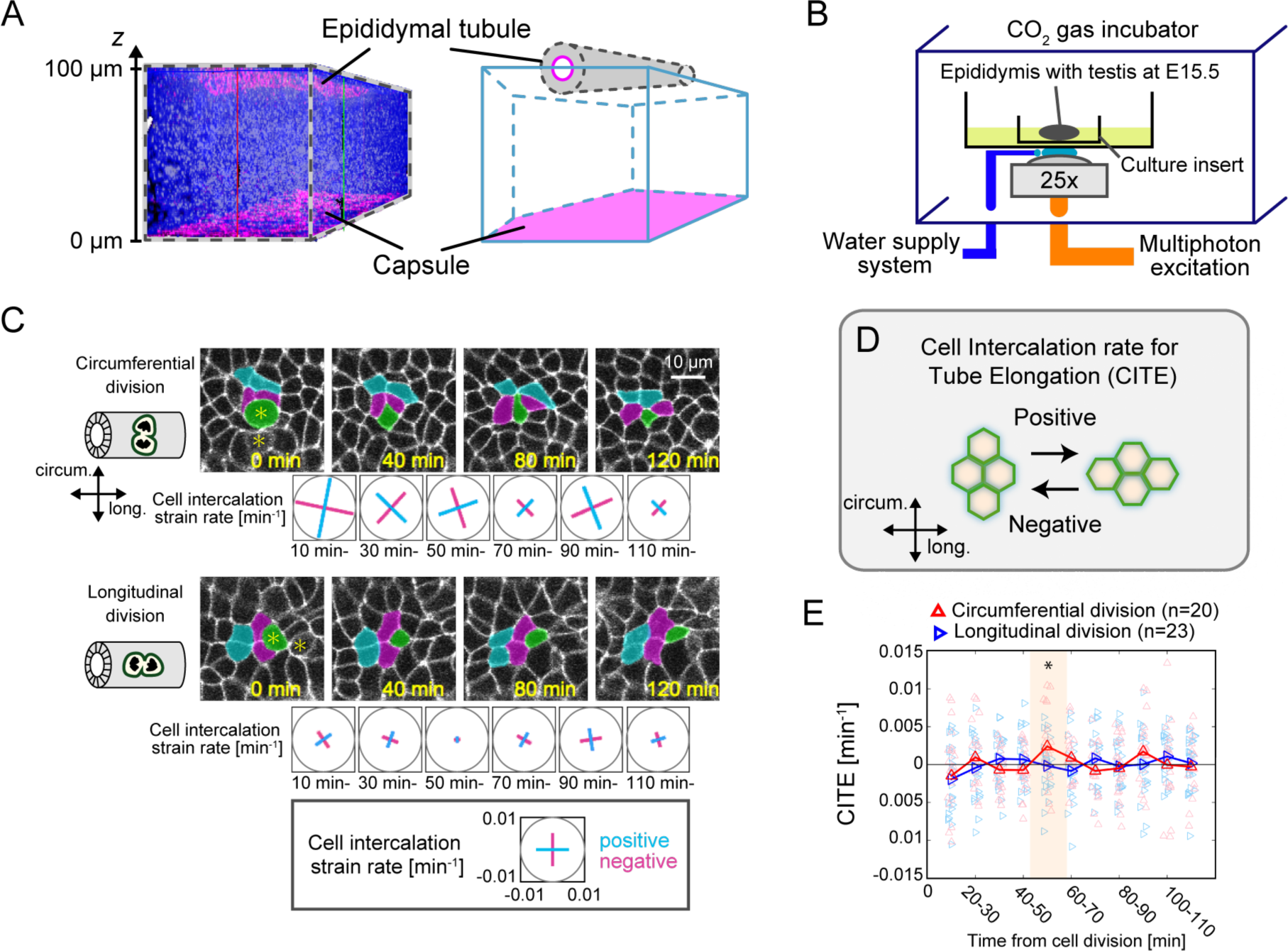
Live imaging revealed that the oriented cell division occurs following to the circumferential cell division. (A) The dimensions of epididymis. The rendering image from the immunofluorescence for the ZO-1 (magenta) (left) and its schematics (right). (B) The schematics for the incubator-integrated multiphoton excitation microscopy system. (C) Snapshots from live imaging data in the case of the circumferential and longitudinal cell divisions. Time origin is at the timing of cytokinesis completion. Asterisks represent daughters of divided cell, one of which is colored artificially in green. Some cells are also colored for easy visualization. The scale and sign for cell intercalation strain rate is shown in the bottom. Scale bar: 10 μm. (D) Schematics for the CITE. (E) The time series data for the case of circumferential cell division (red, *n*=20) and that of the longitudinal division (blue, *n*=23). Time zone in which the mean value is not statistically zero is highlighted.

We focused on the behavior of cells neighboring to mitotic cells on the cell division axis as those cells are more likely to show cellular responses against the mechanical perturbations according to the first suggestion in the previous section. Careful observations led to the discovery of characteristic behaviors in those cells; the oriented cell intercalation for elongating the tubule occurs more frequently against the circumferential cell division compared against the longitudinal one (Fig. 5C and Mov. 2). For quantification of the oriented cell intercalation behavior, we measured the cell intercalation strain rate (Blanchard et al., 2009) for the circumferential ( 0 ≤*θ* ≤*π*/6) or longitudinal cell division ( *π*/3 ≤*θ* ≤*π*/2) (Fig. 5C and S3), and calculated the cell intercalation rate for tube elongation (CITE) as a measure of the oriented cell intercalation in the tubules. Positive/negative values of the CITE indicate that cell intercalations undergo for the tubule elongation/shortening (Fig. 5D; see the Materials and Methods (II-vii) for more details of CITE). From the quantitative analysis, we found that the oriented cell intercalation tends to occur 60 min after the circumferential cell division (*t*-test P<0.05 in the time range: 50-60 min in the case of circumferential cell division) (Fig. 5E).

Considering the cellular movement as an output response, the oriented cell intercalation should be operated through rapid biochemical process such as phosphorylation. In addition to this inference, we demonstrated by the extensive numerical simulations that the cell division within the epithelial tissues could attenuate the cell tension in adjacent cells along the division orientation owing to pressure of the dividing cells to the adjacent ones (Fig. 1F and 4B´). Therefore, we hypothesized that MRLC would be phosphorylated as a cellular response to the reduction of circumferential cell tension provided by the neighboring cells, ultimately driving the oriented cell intercalation.

### Polarized cellular responses to mechanical simulations lead to the maintenance of the tubule radial size

To check whether this hypothesis is rational, we carried out the computer simulation of the vertex dynamics model including a mechano-responsive regime and its counterpart regimes (Mov. 3). We implemented the mechano-responsive regime into the mathematical model as the cells keep constricting one of the circumferential junctions for a parameterized period since the cell tension anisotropy *ρ_α_* reached at the threshold value *ρ\** due to the reception of intense pressure (Fig. 6A and S4; Materials and Methods (III-iv)). The numerical simulation of the model clearly suggests that the mechano-responsive regime can realize spatiotemporally-constant tubes with smaller radial size compared to the case of no regulation (Fig. 6B and D).

**Figure 6.**
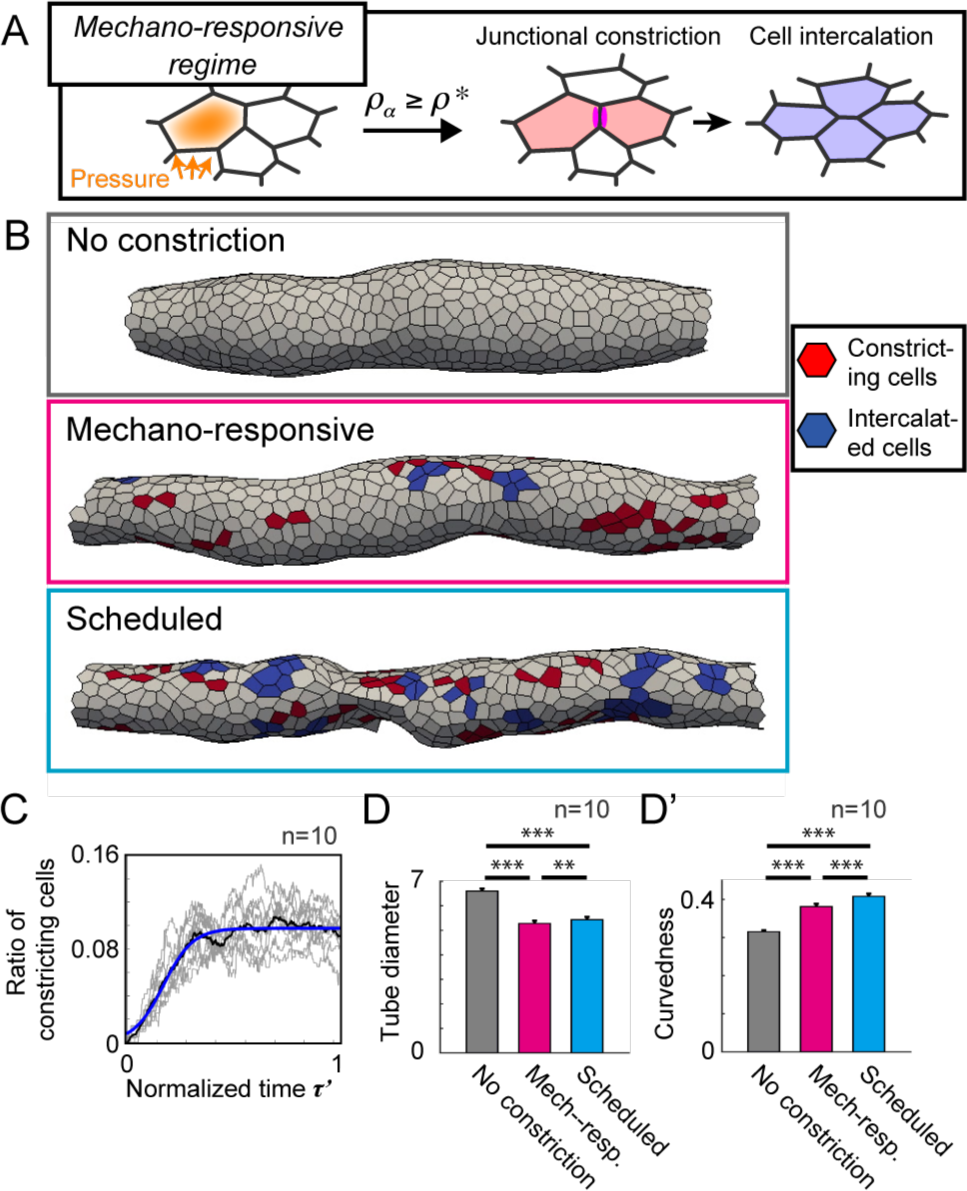
Model simulations suggest that the mechano-responsive cell behavior explains the maintenance of tubule radial size. (A) Schematics for the mechano-responsive regime. In this regime, the circumferential junction constriction starts when the cell tension anisotropy becomes larger than the threshold *ρ*\* due to the pressure from circumferentially-neighboring cells. (B) Generated tubes in different regimes: no constriction, mechano-responsive, and scheduled. Red color in the tubes represents cells constricting the circumferential junction and blue represents cells that have completed the cell intercalation. (C) The ratio of constricting cells to the total cells over the normalized simulation time *τ*\* in the mechano-responsive regime: data in each simulation (gray), the mean value (black) and fitting with the logistic function (blue). *n*=10. (D and D´) Resulting morphological quantities of tubes, the tube diameter and the tube curvedness, in different regimes. See the Materials and Methods (III-vi) for the definition of those morphological quantities. *n*=10.

On the other hand, we performed simulations in a counterpart regime, the 'scheduled’ regime, where the timing for junction constriction was predetermined and the constricting cells were randomly selected. For determining the schedule of the constriction, we sampled theratioofconstrictingcellsalongthenormalizedsimulationtime *τ′* inthe mechano-responsive regime and obtained the function by fitting the mean curve data (Fig. 6C; see Eq. (iv-3) in the Materials and Methods (III-iv)). In the scheduled regime, the tubes can be formed with irregular surfaces and non-constant radial size along the longitudinal axis whereas the radial size was smaller than no regulation as well as the case of mechano-responsive regime (Fig. 6B, D and D´).

In the scheduled regime, the oriented cell intercalation that leads to the local shrinkage of tubules occurs independently to the variations of cell tension, resulting in the irregular surface of tubules. In the mechano-responsive regime, however, the mechanical stress at the supra-cellular scale can be modulated due to the active multicellular movement immediately responding to the mechanical perturbations; therefore, the smooth-curved surface throughout the longitudinal axis of tubules can be realized in this regime. These simulation analyses suggest that the mechano-responsive regime, as an embodiment of our hypothesis, explain the radial size maintenance of proliferating tubes.

### The epididymal tubule cells activate myosin in response to anisotropic mechanical perturbations

To verify the hypothesis, we applied uniaxial compression (~33%) to isolated epididymides embedded in collagen gel using a flexible polydimethylsiloxane (PDMS) chamber, and evaluated the myosin activity as a cellular response to the compression (Fig. 7A). We performed the compression assay for 3 groups, i.e., no uniaxial compression as the control, lateral, and longitudinal compression, to examine the cellular response to different mechanical perturbations (Fig. 7B). After 10 min of continuous load, we quantified the cell shape and configuration altered to the uniaxial compression. For the quantification, we extracted the shape of epididymal cells from the immunofluorescence images for E-cadherin (Fig. 7C), and calculated the aspect ratio and the major axis angle from best-fit ellipses of the cell shapes. The shape and configuration of cells in the uniaxial compression treatments were significantly different from those in the no uniaxial compression, confirming that the epididymal tubule cells were deformed along the compression axis (The Kruskal-Wallis test, *P*<0.001) (Fig. 7D-D´´).

**Figure 7.**
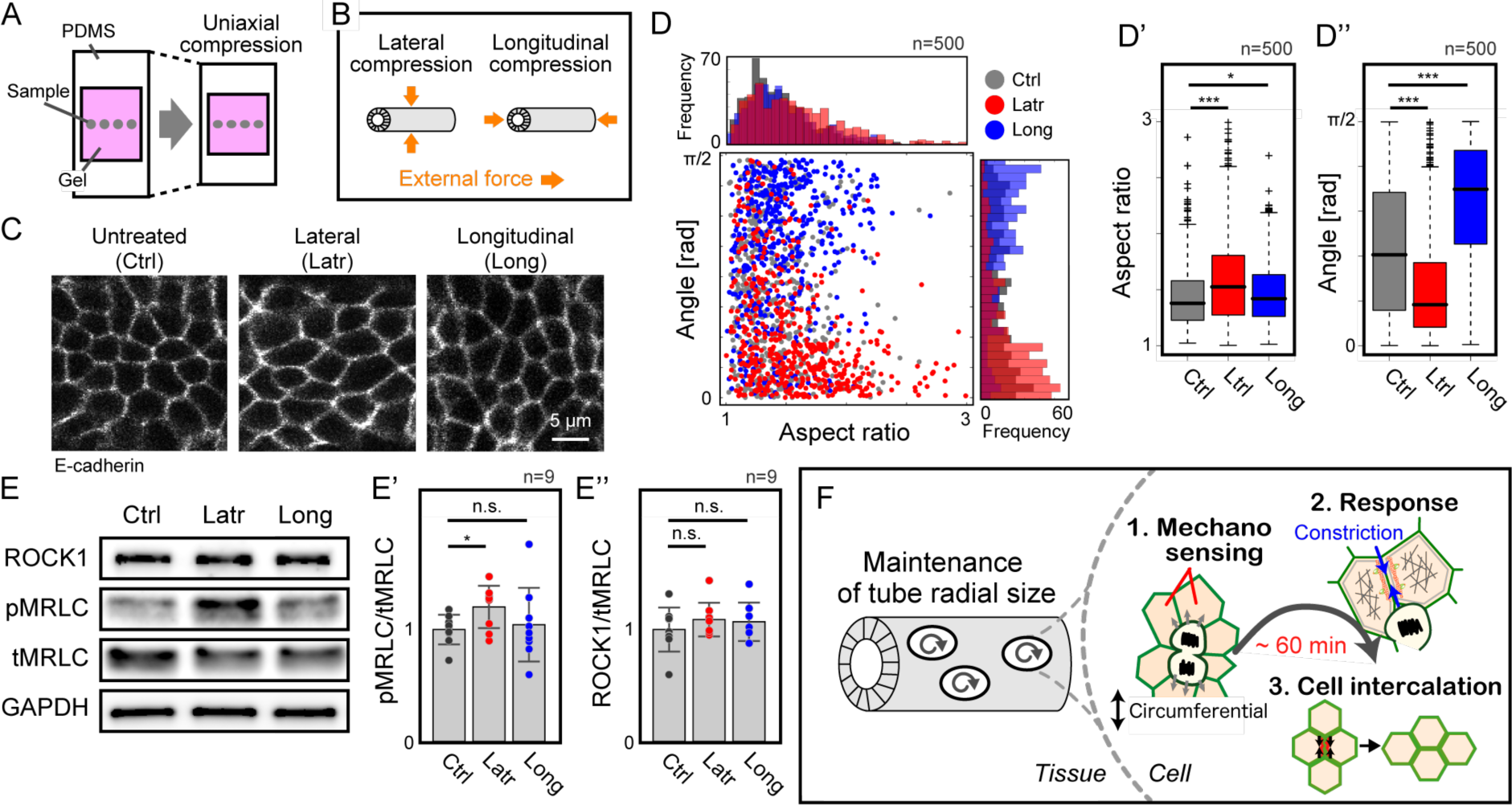
Mechanical perturbation assays uncovered that the myosin is activated anisotropically in the tubule axis. (A and B) Schematics for the uniaxial compression assays. (C) Morphology of the tubule cells following the mechanical treatments. The cell shape was recognized by immunofluorescence for the E-cadherin. Scale bar: 5 μm. (D-D´´) The aspect ratio and the major axis angle for each treatment: control (gray), lateral compression (red), and longitudinal compression (blue). The data are shown as scatter plot and histogram in (D) and as boxplot in (D´ and D´´). *n*=500. (E-E´´) Immunoblotting and quantification of myosin activity and ROCK1 for the mechanical treatments. The tubule cells activate myosin as a cellular response to lateral compression. (E´ and E´´) The data are normalized to the mean value in the control. *n*=9. (F) Model for the maintenance of tubule radial size.

After 20 min, we measured the myosin activity as the relative intensity of pMRLC to the total MRLC (tMRLC) and found that the myosin activity is larger in the lateral compression compared in other cases, indicating that the MRLC phosphorylation is exclusively enhanced due to the lateral compression (The Friedman test, *P*<0.05) (Fig. 7E and E´). Moreover, we quantified the level of MRLC kinase as the ratio of the Rho-associated protein kinase 1 (ROCK1) to the tMRLC, and found that it was not significantly different between treatments (The Friedman test, *P*>0.05) (Fig. 7E and E´´). These results suggest that the epididymal tubule cells possess an intrinsic system in which MRLC phosphorylation is triggered in response to the cell tension reduction along the tubule circumferential axis without changing the total amount of ROCK.

## Discussion

Altogether, we showed that the cells trigger myosin activation leading to oriented cell intercalation by responding to mechanical forces anisotropically in the tubule. Therefore, we propose a system in which the mechanoresponsive behavior of cells at the supra-cellular scale organizes the maintenance of the tubule radial size at the tissue scale (Fig. 4F). In developmental processes, anisotropic growth of proliferating tissues while maintaining an axis, such as tube elongation due to the radial maintenance specifically dealt with in this study, causes generation of compressive forces in the growing axis, eventually leading to a buckling-induced wavy pattern formation (Dong et al., 2014; Hirashima, 2014; Savin et al., 2011). Thus, the proposed polarized mechanoresponse system might work as a regulatory principle underlying mechanisms for self-organized tissue morphogenesis (Beloussov, 2015; Sasai, 2013).

To understand the system further, future work will need to identify what molecular machineries sense the anisotropic mechanical force in tissues and how the received mechanical stimuli are chemically transduced to activate the myosin at apical junctions. It is tempting to speculate that the directional preference of cells would be provided by regulators of planar cell polarity (PCP) that promote the downstream myosin phosphorylation (Bosveld et al., 2012; Guillot and Lecuit, 2013; Nishimura et al., 2012), such as Vang-like (VANGL) and Tyrosine-protein kinase-like 7 (PTK7). These proteins are well aligned circumferentially along junctions of epididymal tubule cells and loss of PCP disrupts the actomyosin-mediated cell intercalation resulting in tubule expansion (Xu et al., 2016) (Fig. S5) as seen in renal tube development (Karner et al., 2009; Kunimoto et al., 2017). Another possibility is that actin filaments may function as polarity tension sensor in cooperation with unknown molecules that specify the circumferential axis because those have been recognized to sense the tension reduction (Galkin et al., 2012; Hayakawa et al., 2011). We believe that our findings move forward to reveal structural components and dynamic properties in the polarized mechanoresponse system that controls tissue morphogenesis.

## Materials and Methods

This section includes (I) Experiments, (II) Quantifications, (III) Mathematical modeling, and (IV) Others (Statistical tests, etc.).

### (I) Experiments

#### (i) Animals

For live imaging analysis, we used fluorescence reporter mice produced by crossing the Pax2-Cre line(Ohyama and Groves, 2004) (a gift from T. Furukawa in Osaka Univ.) and R26R-Lyn-Venus line(Abe et al., 2011). Otherwise, we used imprinting control region (ICR) mice purchased from Japan SLC, Inc. We designated the midnight preceding the plug as embryonic day 0.0 (E0.0), and all mice were sacrificed by cervical dislocation to minimize suffering at E15.5 or E16.5. All the animal experiments were approved by the local Ethical Committee for Animal Experimentation of Institute for Frontier Life and Medical Sciences, Kyoto University, and were performed in compliance with the Guide for the Care and Use of Laboratory Animals at Kyoto University.

#### (ii) Antibodies and Fixative Solutions for Immunostaining

We used the following primary antibodies with dilution ratio 1:200 as a standard condition: rabbit monoclonal anti-E-cadherin (Cell Signaling Technology, #3195), mouse monoclonal anti-γ-Tubulin, (Sigma-Aldrich, #5326), rabbit polyclonal anti-phospho histon H3 (pHH3)(Merck Millipore, #06-570), rat monoclonal anti-pHH3 (Abcam, #ab10543), rabbit polyclonal anti-phosho Myosin light chain (pMRLC) (Abcam, #ab2480), rabbit polyclonal anti-Vangl1 (Atlas Antibody, #HPA025235), goat polyclonal anti-Vangl2 (Santa Cruz Biotechnology, #sc-46561), and mouse monoclonal anti-ZO-1 (Thermo Fischer Scientific, #33-9100). We used the paraformaldehyde (PFA) for anti-E-cadherin, anti-γ-Tubulin, and anti-pHH3; we used the trichloroacetic acid (TCA) for anti-pMRLC, anti-Vangl1, anti-Vangl2, and anti-ZO-1.

#### (iii) Whole-tissue Immunofluorescence and Fluorescent Dye

Dissected epididymides were fixed with 4% PFA in phosphate buffered saline (PBS) for 2 h at 4°C or for 20 min at 37°C, or with 2% TCA in Ca^2+^- and Mg^2+^-free PBS for 20 min at 4°C, depending on target epitopes. The samples were then blocked by incubation in 10% normal goat serum (NGS) (Abcam, #ab156046) or 10% normal donkey serum (Abcam, #ab166643) diluted in 0.1% Triton X-100/PBS, depending on derived animals of secondary antibody, for 3 h at 37°C. The samples were treated with primary antibodies overnight at 4°C, and subsequently incubated with secondary antibodies conjugated to either the AlexaFluor 546 or the AlexaFluor 647 (1:1000, Thermo Fischer Scientific) overnight at 4°C. We used TRITC-conjugated phalloidin (0.3 μg/ml, Merck Millipore, #FAK100) and Hoechst 33342 (5 μg/ml, Thermo Fischer Scientific, #H3570) for the visualization of F-actin and DNA, respectively.

#### (iv) Volumetric Fluorescence Imaging

We mounted samples with 1% agarose gel of 10 μl on a glass-based dish (Greiner Bio-One, #627871) for stable imaging. Then, the samples were immersed with the BABB solution (benzyl-alcohol and benzyl-benzoate, 1:2) or CUBIC2 solution for optical clearing(Hirashima and Adachi, 2015; Susaki et al., 2014; Yokomizo et al., 2012). Finally, we obtained 8-bit volumetric fluorescence images using the confocal laser scanning platform Leica TCS SP8 equipped with the hybrid detector Leica HyD. We used the objective lenses magnifications of ×20 (numerical aperture (NA) = 0.75, working distance (WD) = 680 μm, HC PL APO CS2, Leica), ×40 (NA = 1.3, WD = 240 μm, HC PL APO CS2, Leica), or ×60 (NA = 1.4, WD = 140 μm, HC PL APO CS2, Leica).

#### (v) Explant Organ Culture

The epididymides were surgically excised at a region of the vas deferens close to the tail of epididymis, and those with testis were dissected from the embryonic body. Samples were then placed on a hydrophilic polytetrafluoroethylene organ culture insert with a pore size of 0.4 μm (Merck Millipore, #PICM01250), which was preset in a 35 mm petri dish (Thermo Fischer Scientific, #153066) filled with a culture medium. The culture medium we used was Dulbecco's Modified Eagle's medium (Nacalai Tesque, #08489-45), containing 10% fetal bovine serum (Thermo Fischer Scientific, #12483) and 1% penicillin-streptomycin mixed solution (Nacalai Tesque, #26253-84) at 37°C under 5% CO_2_. The samples were cultured in an air-liquid interface; the total amount of culture medium was 800 μl for the petri dish.

#### (vi) Small-molecule Inhibitors

We used the following inhibitors to suppress or activate the generation of actomyosin-mediated tension: Blebbistatin (Merck Millipore, #203391), Y-27632 (Merck Millipore, #SCM075), and Calyculin A (Cell Signaling Technology, #9902).

#### (vii) Live Imaging

The organ culture setting was according to the condition as described above with the following 2 minor modifications. First, we used a 35 mm glass-based dish (Iwaki, #3910-035) with 700 μl of the culture medium in the dish. Second, we adhered each supporting leg of the organ culture insert on the dish with 20 μl of 1% agarose gel. This preparation prevents the culture insert from sliding during the live imaging. We used an incubator-integrated multiphoton fluorescence microscope (Olympus) using a ×25 water-immersion lens (NA = 1.05, WD = 2mm, Olympus). Imaging conditions were as follows; excitation wavelength: 945nm (Mai-Tai DeepSee eHP, Spectra-Physics), AOTF: 3.5%-1.5%, scan size: 1024 × 320 pixels, scan speed: 10 μsec/pixel, z-axis interval: 1 μm, and time interval: 5 min, magnification: ×2.

#### (viii) Tissue Compression Assay

For the tissue compression assay, we used a manual stretch/compression device (STREX, #STB-10) with a polydimethylsiloxane (PDMS) chamber (STREX, #STB-CH-04), and conducted in the following manner. First, we exposed the PDMS chamber to plasma arc for 1 min using a desktop vacuum plasma processing device (STREX, PC-40) for hydrophilic treatment to enhance the adhesion between the chamber and collagen gel into which the epididymides were embedded. Next, we immediately placed isolated epididymides onto the hydrophilized PDMS chamber in its 50% stretched state, and fulfilled with 500 μl of type I collagen (Nitta Gelatin, Cellmatrix Type I-A), followed by gelation process for 10 min at 37°C. Then, we applied 1 ml of the culture medium into the chamber, and the epididymides with the collagen gel were compressed by relaxing from the 50% stretched state around 0.2 mm/sec. No relaxing the stretched chamber was regarded as control treatment. Finally, the samples were incubated for 10 min for the morphological confirmation analysis and 20 min for the Western blotting analysis at 37°C under 5% CO_2_. In the lateral compression assay, the epididymal tubules were deformed along the circumferential axis in any peripheral region and the circumferential length of individual cells became smaller. We consider this is due to gravity of the culture medium filled on the upper layer of gel as this does not happen without the culture media.

For the immunoblotting assay, we extracted total protein from each treatment, and adjusted the amount to the same among the treatments for the SDS-PAGE as described above. We first confirmed that there were no significant differences of GAPDH and those of MRLC among the treatments. Then, we evaluated the myosin activity and the level of MRLC kinase (ROCK1); even when the ROCK1 level in the epididymides was evaluated as the ratio of ROCK1 to the GAPDH, it showed qualitatively the same results. There was no ROCK2 expression in the epididymides. Finally, the activities were each normalized by mean values of the control group.

#### (ix) Total Protein Extraction and Western Blotting

For total protein extraction, dissected epididymides were immersed in the SDS-free RIPA buffer with protease inhibitor cocktail (Nacalai Tesque, #08714) supplemented with the EDTA-free phosphatase inhibitor cocktail (1:100, Nacalai Tesque, #07575), and were disrupted by an ultrasonic cell disruptor (Microson). The lysates were placed onto the ice for 20 min and were centrifuged at 13,000 ×g for 10 min at 4°C. Protein concentration of the supernatant was determined by the bicinchoninic acid assay.

The lysates were prepared for SDS-PAGE by adding 2×Laemmli sample buffer (Bio-Rad, #161-0737) with 2-mercaptoethanol (Bio-Rad, #161-0710) and by boiling at 96°C for 5 min. Next, the lysates containing approximately 5 μg of proteins were loaded into each lane of the Mini-PROTEAN precast gels (Bio-Rad, #4569035), and electrophoresis was carried out in Tris/glycine/SDS running buffer (Bio-Rad, #1610732) at constant 150V for 35 min. Then, the proteins were blotted onto 0.2 μm polyvinylidene difluoride membrane (Bio-Rad, #1704272) in HIGH MW mode (1.3A, 25V for 10 min) of the Trans-Blot^®^ Turbo^TM^ Transfer System (Bio-Rad, #170-4155) for ROCK1 detection and in the LOW MW mode (1.3A, 25V for 5 min) for others.

The blotted membranes were then immersed in 15% H_2_O_2_/Tris-buffered saline (TBS) solution for 30 min at room temperature for blocking endogenous peroxidase followed by blocking with 5% NGS at 37°C for 60 min. For immunoblotting, the membranes were incubated with primary antibodies diluted in 0.1% TBS/Tween-20 at 4°C over night. The concentrations of antibodies used were 1:100,000 for mouse monoclonal anti-GAPDH (Wako, #015-25473), 1:500 for rabbit polyclonal anti-myosin light chain 2 (Cell Signaling, #3672) and mouse monoclonal anti-phospho-myosin light chain 2 (Cell Signaling, #3675), and 1:2000 for rabbit monoclonal anti-ROCK1 (Abcam, #ab45171). Then, the membranes were incubated with diluted secondary antibody solutions in TBS/Tween-20 at 37°C for 60 min; the concentration was 1:50,000 for goat anti-rabbit IgG conjugatd to horseradish peroxidase (HRP) (Santa Cruz Biotechnology, sc-2004) and goat anti-mouse IgG conjugated to HRP (Santa Cruz Biotechnology, sc-2005). Finally, protein bands were detected using the Amersham ECL Select Western Blotting Detection Reagent (GE Healthcare, #RPN2235), and were scanned with ImageQuant LAS 4000 (GE Healthcare).

### (II) Quantifications

#### (i) Cell Division Orientation

We first performed coimmunolabeling for γ-Tubulin and pHH3 of 6 different epididymides to detect microtubule organizing centers of mitotic cells as described in this section (I)-(iii) with a minor modification. That is, the samples were incubated in 10 mM sodium citrate buffer including 0.1% Triton X-100/PBS for 20 min at 80°C for heat-induced retrieval of γ-Tubulin epitope before blocking treatment. Next, we obtained volumetric images of 1024 × 256 pixels as described above using ×40 lens with z-axis interval of 1 μm. Third, we manually measured three angles *α*_1_, *α*_2_, and *α*_3_ shown in Fig. 2B for each mitotic cell to define a local coordinate system (*O*’; see Fig. 2C), origin of which corresponds to the center position of mitotic cell. Each orthogonal basis of *O*’-coordinate is defined as follows: longitudinal direction of epididymal tubule (*x*’), surface normal of epididymal tubule (*z*’), and orthogonal direction for both *x*’ and *z*’ according to the right-handed system (*y*’). Then, we manually measured positions of the two γ-Tubulin-positive dots (microtubule organizing centers: MTOCs) in a pHH3-positive cell, and obtained two angles *φ* and *θ* in a local sphere coordinate (Fig. 2D) resulted from coordinate transformation into the *O*’-system. Finally, we selected samples under a criterion that the distance between the two γ-Tubulin-positive dots is more than 1.5 μm. We regarded the samples whose *φ* is less than 40 degree as cells dividing in parallel to the surface of epididymal tubule.

#### (ii) Apical Junction Morphology and pMRLC/VANGL Signal Intensity

We performed coimmunostainining for ZO-1 and either pMRLC or VANGL2 of 8 different epididymides as described above with a minor modification: the dilution ratio of anti-pMRLC was 1:400 for preventing saturation binding. We obtained volumetric images of 1024 × 256 as described above using ×60 lens with z-axis interval of 0.3 μm. Then, we separated the images into smaller ones around 200 pixels (≈36 μm) on a side so that the slope of epididymal tubule can be regarded as a linear line.The longitudinal The digital image processing was performed as follows. First, images for ZO-1 were filtered slice by slice with a 2D median filter (3 × 3 pixels) followed by a 3D Gaussian filter with a standard deviation of 3 pixels to reduce undesired noise. Next, the filtered images were binarized using the 3D maximum entropy thresholding method (Kapur et al., 1985), and the maximum one among the binarized component of 26-connected pixels was regarded as an apical junction network. Then, we performed 3D skeletonization process for the images of apical junctions, and obtained a graph network including nodes and edges of skeletonized voxels utilizing distributed MATLAB codes with modifications (Kerschnitzki et al., 2013). Finally, to extract the skeleton of apical cellular junctions (Fig. 3A), we screened out the skeletonized edges to avoid over-skeletonization results in the following 3 criteria; (1) The edge length between two nodes is less than 10 μm, (2) the edge length is shorter than two-fold linear length between two nodes linked with the corresponding edge, (3) the number of edges from a node is more than 3. The screening process under the criteria produced 96% left from the total number of raw skeletonized edges. The relative angle of apical junction edges against the longitudinal axis of epididymal tubule was calculated as 
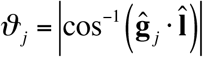
 where *j* is an index of apical edges, **ĝ** is an unit vector of apical edges, and **Ȋ** is that of the longitudinal axis of tubule, as shown in Figure 3B. For the signal quantification, we extracted signal intensity of pMRLC or that of VANGL2 on the apical cellular junctions by using a thickened skeleton of apical junctions with a dilation operator (3 × 3 × 3 voxels), and calculated average values for each edge of thickened apical junction skeletons.

#### (iii) Tubule Diameter in Inhibitor Assay

First, we performed immunostaining for the E-cadherin of 8 different epididymides for each inhibitor treatment to visualize the epididymal tubule as described above, and obtained the volumetric images of 512 × 512 pixels using ×20 lens with z-axis interval of 5 μm. We utilized the built-in module of a software platform, the Leica Suite X, (Leica), to create a montage image of whole epididymis. Then, we manually extracted epididymal tubules from maximum intensity projection images by erasing the efferent ductules and noise pixels due to non-specific staining. From the preprocessed images, we obtained the centerline of epididymal tubule by applying the built-in skeletonization algorithm in MATLAB for the binary images obtained with arbitrarily determined thresholds, and finally calculated the diameter along the centerline of tubule.

#### (iv) pMRLC Signal Intensity along Epididymal Tubule

First, we performed immunostaining for the pMRLC of 12 different epididymides as described above with a minor modification: the dilution ratio of anti-pMRLC was 1:100. Second, we obtained the volumetric images of 512 × 512 as described above using ×20 lens with z-axis interval of 1 μm; we manually made montage images to avoid an overlap region of composite images. Third, we manually erased signal intensity on pixels corresponding to non-apical cellular region from the MIP images to obtain the centerline of epididymal tubule as described above. Then, we calculated average values of pMRLC intensity within a circle with a radius of 9 um along the epididymal tubule. Finally, we obtained the pMRLC signal distribution along the epididymal tubule by smoothing the averaged signal intensity with a moving average filter with a span of 40 μm – data in 20 μm range from the both ends of tubule were omitted. The maximum value of vertical axis in Fig. 3H is 1 which corresponds to the maximum intensity of 8 bit images. The horizontal axis of Fig. 3H was normalized by the tubule length of each sample.

#### (v) Correlation between the Number of Mitotic Cells and pMRLC Signal Intensity

We used the data on the number of mitotic cells from our previous work (Hirashima and Adachi, 2015) and one on the pMRLC intensity obtained in this study. We divided 30 bins along the epididymal tubule and calculated maximum likelihood estimates with the Poisson distribution for the number of mitotic cells and with the Gaussian distribution for the pMRLC intensities in each bin. The Pearson’s correlation coefficients are 0.49 (*P*-value=0.006) for E15.5 and 0.56 (*P*-value=0.001) for E16.5.

#### (vi) Apical Surface Area

We obtained 1024 × 256 volumetric images of fluorescently-labeled F-actin in epididymal tubules as described above using ×40 lens with z-axis interval of 1 μm, and randomly collected samples of apical cell surface from image planes of 3 different epididymal volumetric data on which the apical cell surfaces exhibited. Then, we calculated the apical surface area by tracing the peripheral of apical cell surfaces manually.

#### (vii) Cell Intercalation Rate for Tube Elongation (CITE)

We first obtained time-lapsed volumetric images for epithelial dynamics from the head region of 4 different samples at E15.5 through the live imaging as described above, and collected time series data for a group of cells around mitotic cells. In particular, we sampled those cells on an image plane perpendicular to the optical axis from the image data, and focused on cross sections in-between the apical and basal part of cells. We defined cytokinesis as an origin of time for our analysis, and analyzed data on cell intercalation dynamics with time interval of 10 min ( Δ*t* = 10 min).

Next, based on an earlier study, we calculated the cell intercalation strain rate tensor 
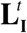
 at time *t* as follows:

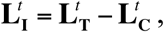

where 
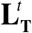
 is a local tissue velocity gradient tensor, and 
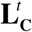
 is a cell shape strain rate tensor (Blanchard et al., 2009). Because our analysis was performed in planar, the tensors are 2 × 2 matrices. For the calculation of a local tissue velocity gradient tensor 
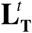
, we specified one or two neighboring cells of a daughter cell from a mitotic cell that located an extension of the division axis as central cells, and marked the adjacent cells of the central cells including the central cells and the daughter cell as domain cells. Then, we manually traced each one of domain cells to obtain cell areas, cell shapes with the best fit ellipses, and cell centroids 
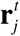
, where *j* is an index of domain cells (see marked cells by yellow lines in Fig. S4A). Using data of the cell centroids 
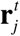
 and that of velocities 
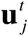
, the local velocity gradient tensor was obtained by the least squares approximations according to the following equation,

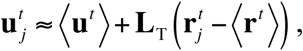

where 
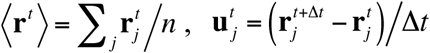
 and 
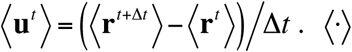
 indicates an average within the domain.

For the calculation of the cell shape strain rate tensor 
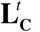
, we used lengths of principal axes of a fitted ellipse and angle of the major axis from the longitudinal axis of epididymal tubule to obtain a matrix 
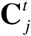
 that represents the cell ellipse. Then, we calculated 
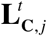
 from 
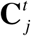
 and 
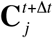
, and 
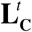
 was calculated by taking the area-weighted average of 
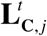
 within the domain cells. Note that we normalized 
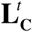
 to satisfy that a trace of 
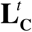
 was equal to that of 
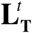
, realizing that the cell intercalation strain rate tensor 
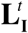
 had zero dilation.

Finally, we obtained the CITE at time 
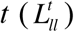
 from an element of the cell intercalation strain rate tensor:

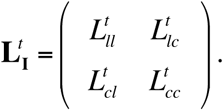

Note that the 
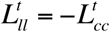
.

The data were classified into two groups depending on the relative angle of cell division axis to the longitudinal axis of epididymal tubule – we defined the relative division angle of 0-30 degrees as longitudinal division and that of 60-90 degrees as circumferential division.

### (III) Mathematical Modeling

#### (i) Basic Formalization of the Vertex Dynamics Model

We employed a vertex dynamics model (VDM), a type of cell-oriented model, to represent multi-cellular dynamics in epithelial tissues (Farhadifar et al., 2007; Fletcher et al., 2014; Nagai and Honda, 2001; Rauzi et al., 2008). In this model, a cell is geometrically regarded as a polygon/polyhedron, of which vertices are elementary points that constitute the cell shape, and a group of cells can be represented by a set of polygons/polyhedrons shared by neighboring cells (Fletcher et al., 2013; Honda and Eguchi, 1980; Honda et al., 2004; Nagai and Honda, 2001; Okuda et al., 2013). In this study, we paid notice to apical junctions of the epididymal tubule on which a marker of tension generator pMRLC is only localized (Fig. 2A), and modeled the developing single-layered epithelial tube as a flexible multicellular membrane in 3D space according to earlier studies (Du et al., 2014; Osterfield et al., 2013).

In the VDM, the dynamics of position of vertex *i*, **r**_*i*_, obey the equation of motion based on the principle of least potential energy *U* as follows:

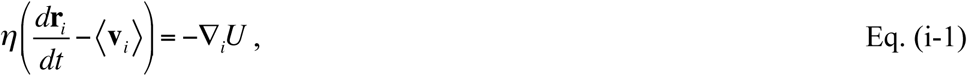

where *η* is a viscosity coefficient, and 〈**v***^i^*〉 is a local velocity of vertex *i*, defined as

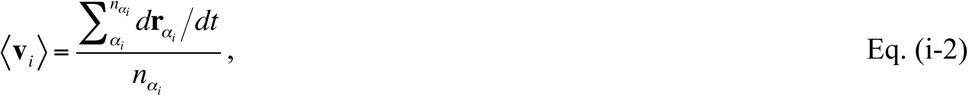

where *α_i_* is an index for cells contacting to vertex *n_α_i__* is a number of the cells contacting to vertex *i*, and **r***_α_i__* is a centroid of cell *α_i_*(Mao et al., 2013; Okuda et al., 2014). For a potential energy as a minimum expression to represent epididymal tubule cells, we defined as

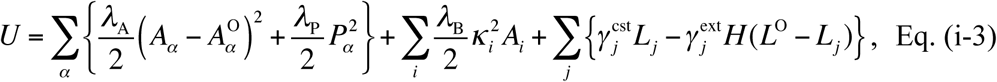

according to earlier studies (Collinet et al., 2015; Farhadifar et al., 2007; Rauzi et al., 2008). *α*, *i*, and *j* each denote an index of cells, that of vertices, and that of junctions/edges. The first term represents the cell elasticity; *λ*_A_ is its coefficient, *A_α_* is area of cell *α*, and 
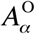
 is a target area of cells. The second term represents isotropic actomyosin contractility at the periphery of apical junctions; *λ*_P_ is its coefficient and *P_α_* is perimeter of the cell *α*. The third term represents bending energy of the flexible epithelial membrane embedded in 3D space; *λ*_B_ is its coefficient, *κ*_i_ is a discrete mean curvature defined at vertex *i* of triangular meshes (Meyer et al., 2003), and *A_i_* is summation of an area of triangular mesh fractions around vertex *i*, each explained one by one. For calculation of the discrete mean curvature at a vertex, let surface of the modeled epithelial membrane be a set of triangular meshes consisting of both the vertex of cells and the centroid of cells (Fig. S1A). Then, we defined the discrete mean curvature at vertex *i* as

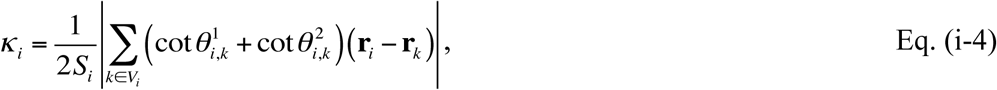

where *k* is an index of the triangular elements included in the set of 1-ring neighbor vertices of the vertex *i*, *V_i_*, *S_i_* is total area of triangular meshes in *V_i_*, 
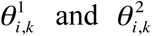
 are each angle depicted in Figure S1A’. See (Meyer et al., 2003) for more details. For calculation of *A_i_*, let us consider triangles consisting a vertex and its 1-ring neighbor cell centroids illustrated in Figure S1A”, and then sum the all triangles up around the vertex. In earlier studies, bending energy was defined as a summation of inner product of unit normal vectors to the surfaces between the adjacent cells (Du et al., 2014; Osterfield et al., 2013). This is valid when all mesh size is ideally equal (Kantor and Nelson, 1987; Seung and Nelson, 1988); therefore, we introduced the bending energy function based on the discrete curvature. The last term represents anisotropic junction constriction/extension; 
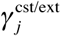
 is its coefficient assigned to each edge, *L_j_* is length of edge, *L^O^* is a target edge length, and *H* is the Heaviside step function. For the simulation in Fig. 1F-F”, 4, and S1, 
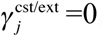
. The details of this term are described later.

#### (ii) Normalized form of the model

By introducing a characteristic cell size*A*_0_, the equations were transformed into nondimensionalized form to reduce the number of parameters as

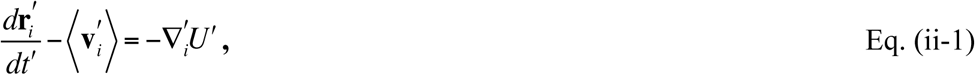

where 
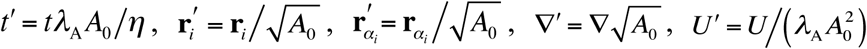
, and

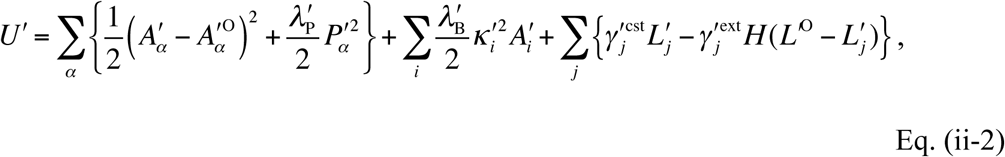

where 
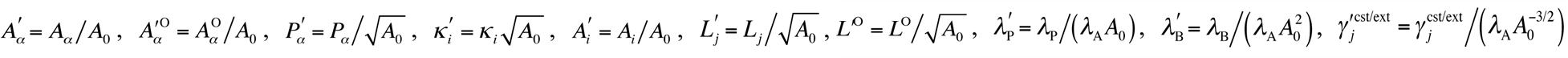
. Inaddition, we introduced a normalized simulation time defined as *τ*′ = *t*′/*T*^cycle^, where *T*^cycle^ is an average cell cycle length, which will be explained. We used these variables and equations, Eq. (ii-1) and (ii-2), for the numerical simulations in this study.

#### (iii) Cell intercalation, Proliferation, and Simulation Conditions

As for the cell intercalation, we assumed that edge rearrangement occurred when the normalized length of edge became smaller than 0.05 and the edge length was set to be 0.1 after the edge rearrangement. See (Fletcher et al., 2013) for details of the edge rearrangement.

As for the cell proliferation, we introduced the average cell cycle length *T*^cycle^, and set cell cycle length of each cell 
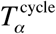
 obey the normal distribution of which the mean is *T*^cycle^ and the standard deviation is 5% of the mean, i.e., 
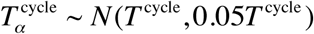
. Also, we introduced normalized cell cycle time of each cell as 
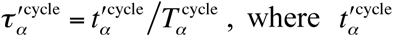
 is nondimensionalized cell cycle time of cell *α*. 
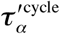
 at *t*′ =0 was assigned within 0 to 0.95 in uniform distribution. As the average cell cycle length in the epididymal tubule can be roughly estimated as 24 hours and cell volume tends to get increasing within the apical surface of epididymal tubule since 50-100 min before the cytokinesis, we simply modeled the cell dynamics during cell cycle as 
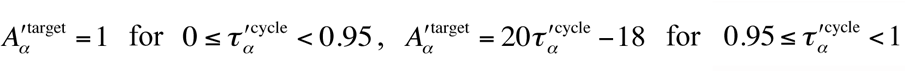
, and the cell divides with returning 
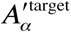
 of daughter cells to 1.The type of cell division orientation was set as follows – circumferential/longitudinal division: uniform distribution in a range of ±0.05*π* from the circumferential/longitudinal axis of tube, and non-biased division: uniform distribution in all range of *θ* defined in the local coordinate at cell (Fig. 1C).

As initial condition for the simulation, we set that a virtual tube was composed of 14 cells along the circumferential direction based on our observation and of 30 cells along the longitudinal axis as for extensive numerical simulations. The cross-sectional center of the tube was set at *y*=0 and *z*=0 of global coordinate and the tube was put along the *x* axis. As boundary condition, the following energy was added to the original potential energy *U*′ at tip cells of the tube:

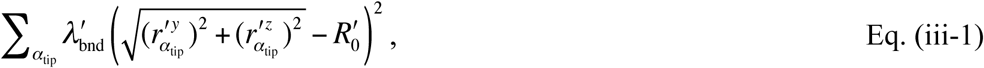

where *α_tip_* is an index for the tip cells of tube, 
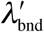
 is its coefficient, 
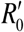
 is a normalized mean radius of the tube to a characteristic cell length 
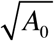
 in the initial setting, 
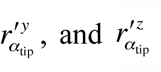
 each represent the relative position of center of cells *α_tip_* on *y* and *z* axis to 
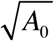
. This energy restricts displacement of the cells *α^tip^* in radial direction of the tube, which imitates physical constraints by the vas deferens and the efferent ducts as boundary tissues of the epididymal tubule. We set 
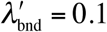
 in the simulations. The results do not vary qualitatively even without the boundary energy term Eq. (iii-1).

#### (iv) Mechano-responsive and Scheduled regimes

In the following, we explain the last two terms of Eq. (ii-2): the anisotropic edge constriction and extension after the cell intercalation. We here introduce a state variable assigned to each cell, Θ*_α_* = {0,1, 2} ; Θ*_α_* = 0,1, 2 represent a naïve state (colored in white, Fig. 6B), edge-constriction state (red, Fig. 6B), and edge-extension state (blue, Fig. 6B), respectively. For all cells, Θ*_α_* = 0, 
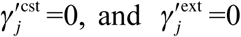
,unless otherwise noted. In our simulation, the following procedures were conducted for both 'mechano-responsive’ and 'scheduled’ regime.

First, cells *α*_′_ that might become a constricting cell were selected and let the state variable of the cell Θ*_α′_* =  1 ; the way of selecting cells depends on each regime, i.e, the mechano-responsive or the scheduled regime. Second, one of the edges that orients to circumferential axis of the tube (the angle *ϑ_j_* is within 70° – 90°) was selected among all edges composing the cells *α*′, and index the edge *j*′.Then, state of cells including the edge *j*′ turned to *Θ_α′_* = 1. Third, the constriction of edge *j*′ was performed according to a schedule as shown in Figure S4A; we set the edge kept constricting in a period *τ^′cst^* via transition period *τ^′trns^* from the starting point of edge constriction by an assumption with phosphorylation process of pMRLC. The maximum value of 
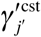
 was set to be *γ^′cst^*. Θ_*α*′_ and 
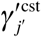
 return to 0 at the end of edge constriction process. In this study, parameter dependence of *γ^′cst^* and that of *γ^′cst^* were examined and determined (Fig. S4B), and we set *γ^′trns^*  0.02. Finally, if the edge arrangement occurred during the process of edge constriction ( Θ_*α*′_ = 1), then the state of cells having the rearranged edge changes to Θ_*α*′_ = 2 and the edge becomes an extension mode, in which the rearranged edge converges to a constant target length; 
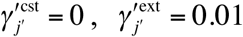
, and *L*^′O^ = 0.1. We set the period of edge extension as *τ^′ext^* = 0.02. Θ_*α*′_ and 
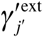
 return to 0 after the process of edge extension. Our implementation for the edge extension process is based on an earlier report that the junction after the cell intercalation grows due to local polarized force driven by medial actomyosin contraction (Collinet et al., 2015).

As for the way of selecting the cells *α*′ in the first step described above, we adopted the two: mechano-responsive and scheduled regime. On the mechano-responsive regime, we supposed that the epididymal tubule cells could receive mechanical stress provided by adjacent cells, and a stress tensor in cells *α* in the local cell coordinate system *O’* was defined as

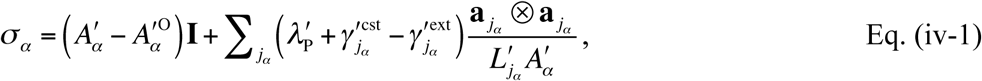

where **I** is a unit tensor, *j_α_* is an index of edges composing the cell *α*, **a** is a vector representing the relative position of two vertices composing the edge *j_α_* (Batchelor, 1970; Ishihara and Sugimura, 2012; Sugimura and Ishihara, 2013). Denoting a diagonal element of the tensor *σ_α_* corresponding to the longitudinal axis of the tube and that to the circumferential axis as 
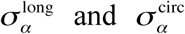
, we then defined the cell tension anisotropy as

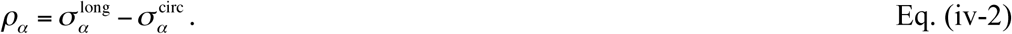

Notethatthecelltensionanisotropy *ρ_α_* is a time-dependentvariable.Inthe mechano-responsive regime, the cells were assigned as *α′* when *ρ_α_* was over a threshold *ρ*\*, of which parameter dependency for the resulting tube morphology was examined (Fig.S4B).

On the scheduled regime, the cells *α′* were randomly selected but timing of the selection was predetermined according to the schedule shown in Figure 6C; Figure 6C shows the fraction of cells having constricting edges to all tube cells in the mechano-responsive regime (*n*=10, gray), and its mean (black). In our simulation, we used the following logistic function (blue) by fitting the mean curve with the least-squared method:

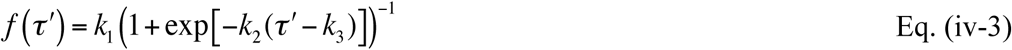

with *k_1_* = 0.1, *k_2_* = 0.01, *k_3_* = 0.17.

#### (v) Morphological and Mechanical Quantities in Simulation

We used the following morphological quantities in the simulation; 1) tube length, i.e., longitudinal linear length between distal edges of tube, 2) tube diameter, i.e., mean diameter of circumferentially-averaged diameter through the centerline of tube, and 3) curvedness, i.e., averaged curvature of the vertices throughout the tube defined as 
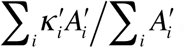
.

For the cell tension anisotropy *ρ_α_*, refer to the Eq. (iv-2).For the tube tension anisotropy in Figure 1F, the stress tensor in the tube was defined as

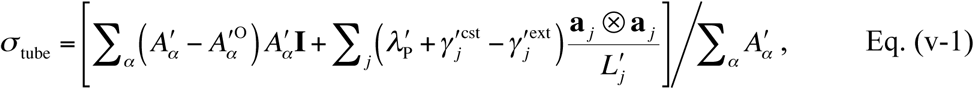

according to earlier studies (Batchelor, 1970; Ishihara and Sugimura, 2012), and using its diagonal element 
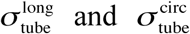
, we introduced

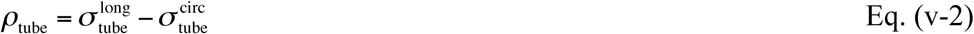

as the tube tension anisotropy. 
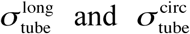
 each correspond to the tube longitudinal tension and the tube circumferential tension appeared in Figure S1.

#### (vi) Determination of Parameter Values (Fig. S1 and Fig. S4)

Parameter values used in this study were (i) 
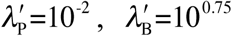
, (ii) *T*^cycle^=10^3.75^, (iii) *ρ*\* = 0.1, *γ*^′cst^ = 1, and *τ*^′cst^ = 0.167 unless otherwise noted. As for (i), we determined the values, at which there were less variations at *t*′=1000 from the initial configuration (Fig. S1B and S1B’). Note that no proliferation was implemented for this analysis. As for (ii), we examined the parameter dependence of cell proliferation rate 1/*T*^cycle^ to the morphological and the mechanical quantities at *τ*′ = 1 (Fig. S1C). The results show that the cell division orientation less affected to the tube shape (length and diameter) in faster cell proliferation rate. In addition, smooth surface of tube was not maintained if the cell proliferation rate was larger than 10^−3.5^ (see curvedness in Fig. S1C). This is because the cell proliferation rate is faster than relaxation time of tube in the dynamics. We determined as *T*^cycle^=10^3.75^. As for (iii), we examined the parameter dependence of *ρ*\*, *γ*^′cst^, and *τ*^′cst^ to the tube shape and its variance at *τ*′ = 1 (Fig. S4B). We first determined *ρ*\* = 0.1 because the coefficient of variation (CV) became larger at *ρ*\* = 0.08 and there were less differences of the mean values to the variation of *γ*^′cst^ at *ρ*\* = 0.125. Then, we determined *γ*^′cst^ = 1 because this point is at a boundary for both the mean and the CV. Finally, as for *τ*^′cst^, we arbitrarily set the value because there were less differences in any values.

### (IV) Others

#### (i) Statistical Hypothesis Testing

Exact information of the statistical tests, sample sizes, test statistics, and *P*-values were described in the Supplemental Information. Statistical hypothesis testing was performed according to (Zar, 2014). *P*-values of less than 0.05 were considered to be statistically significant in two-tailed tests, and were classified as 4 categories; * (*P*<0.05), ** (*P*<0.01), *** (*P*<0.001), and n.s. (not significant, i.e., *P* ≥ 0.05).

#### (ii) Replicates and Sample Number

All replicates in this study are biological replicates. The *n* in the figure legends indicates the number of samples used for each analysis as described below.

Figure 1D and E: *n* indicates the number of cells sampled from 6 epididymides collected from a pregnant mouse.

Figure 2D-F´: *n* indicates the number of apical edges sampled from 8 epididymides collected from 2 pregnant mice.

Figure 2G´: *n* indicates the number of measured points sampled from 8 epididymides collected from 3 pregnant mice.

Figure 3B and B´: *n* indicates the number of epididymides collected from 3 pregnant mice.

Figure 3D´ and D´´: *n* indicates the number of cells randomly chosen from 3 epididymides collected from a pregnant mouse.

Figure 5E: *n* indicates the number of a group of cells sampled from 4 epididymides each collected from 4 pregnant mice.

Figure 7D-D´´: *n* indicates the number of cells randomly chosen from 10 epididymides collected from 3 pregnant mice.

Figure 7E´ and E´´: *n* indicates the number of pregnant mice.

#### (iii) Graph

Except for boxplot, we used mean as representative values, and error bars represent standard deviation. Boxplots were drawn using R (GNU project).

#### (iv) Software

For digital image processing, we used MATLAB (MathWorks) and Image J (National Institute of Health). For graphics, we used MATLAB (MathWorks), R (GNU project), Gnuplot (Free Software), and Paraview (Kitware). For statistical analysis, we used MATLAB (MathWorks) and R (GNU project).

## Additional files

Additional files include full information of statistical hypothesis testing, and supplementary five figures.

## Author Contributions

T.H. performed experiments and simulations, and analyzed all data; T.H. and T.A. wrote the manuscript.

## Acknowledgements

This work was supported by the Platform Project for Supporting in Drug Discovery and Life Science Research (Platform for Dynamic Approaches to Living System) from the MEXT and the AMED, and by the JSPS KAKENHI grant 15K18541. We would like to thank T. Furukawa of Osaka Univ. for kindly providing the Pax2-Cre mice.

## Statistical Hypothesis Testing

The statistical tests, sample sizes, test statistics, and *P*-values were described below. *P*-values of less than 0.05 were considered to be statistically significant in two-tailed tests, and were classified as 4 categories; * (*P*<0.05), ** (*P*<0.01), *** (*P*<0.001), and n.s. (not significant, i.e., *P* ≥ 0.05).

### ›Figure 1D, *φ* at E15.5

Note: The range of data was transformed to 0-360º for circular distributions.

*H*_0_: The population of *φ* at E15.5 is uniformly distributed around the circle.

*H_A_*: The population of *φ* at E15.5 is not uniformly distributed around the circle.

The Rayleigh test. *n*=277, Test statistic: *Z*=35.51, *P*<0.001; Reject *H*_0_.

### ›Figure 1D, *θ* at E15.5

Note: The range of data was transformed to 0-360º for circular distributions.

*H*_0_: The population of *θ* at E15.5 is uniformly distributed around the circle.

*H_A_*: The population of *θ* at E15.5 is not uniformly distributed around the circle.

The Rayleigh test. *n*=197 (0-40º), Test statistic: *Z*=0.40, *P*=0.67; Do not reject *H*_0_.

The Rayleigh test. *n*=277 (0-90º), Test statistic: *Z*=0.25, *P*=0.78; Do not reject *H*_0_.

### ›Figure 1D, *φ* at E16.5

Note: The range of data was transformed to 0-360º for circular distributions.

*H*_0_: The population of *φ* at E16.5 is uniformly distributed around the circle.

*H_A_*: The population of *φ* at E16.5 is not uniformly distributed around the circle.

The Rayleigh test. *n*=358, Test statistic: *Z*=27.32, *P*<0.001; Reject *H*_0_.

### ›Figure 1D, *θ* at E16.5

Note: The range of data was transformed to 0-360º for circular distributions.

*H*_0_: The population of *θ* at E16.5 is uniformly distributed around the circle.

*H_A_*: The population of *θ* at E16.5 is not uniformly distributed around the circle.

The Rayleigh test. *n*=240 (0-40º), Test statistic: *Z*=1.95, *P*=0.14; Do not reject *H*_0_.

The Rayleigh test. *n*=358 (0-90º), Test statistic: *Z*=0.57, *P*=0.56; Do not reject *H*_0_.

### ›Figure 2E´

Note: The data were transformed on logarithmic scale.

*H*_0_: The mean intensities of pMRLC among the three groups of junction angle are all equal.

*H_A_*: The mean intensities of pMRLC among the three groups of junction angle are not all equal.

One-way analysis of variance. *n*=6,259 (Longitudinal), *n*=8,457 (Intermediate), *n*=7,323

(Circumferential), Test statistic: *F*=165.47, *P*<0.001; Reject *H*_0_.

Multiple comparisons using the Tukey-Kramer Test with unequal sample sizes: *P*<0.001 (Long. vs. Intm.), *P*<0.001 (Long. vs. Circ.), and *P*<0.001 (Intm. vs. Circ.).

### ›Figure 2F´

Note: The data were transformed on logarithmic scale.

*H*_0_: The mean intensities of pMRLC among the three groups of junction length are all equal.

*H_A_*: The mean intensities of pMRLC among the three groups of junction length are not all equal.

One-way analysis of variance. *n*=8,691 (Short), *n*=8,749 (Middle), *n*=3,719 (Long), Test statistic: *F*=306.65, *P*<0.001; Reject *H*_0_.

Multiple comparisons using the Tukey-Kramer test with unequal sample sizes: *P*<0.001(Short. vs. Middle), *P*<0.001 (Short vs. Long), and *P*<0.001 (Middle vs. Long).

### ›Figure 3D´

*H*_0_: The circumferential edge length is the same in all three regions of epididymal tubule.

*H_A_*: The circumferential edge length is not the same in all three regions of epididymal tubule.

The Kruskal-Wallis test, *n*=200 for all groups, Test statistic: *H*=14.84, *P*<0.001; Reject *H*_0_.

Steel-Dwass multiple comparisons: *P*=0.016 (Head vs. Body), *P*=0.001 (Head vs. Tail), *P*=0.61 (Body vs. Tail).

### ›Figure 3D´´

*H*_0_: The apical surface area is the same in all three regions of epididymal tubule.

*H_A_*: The apical surface area is not the same in all three regions of epididymal tubule.

The Kruskal-Wallis test, *n*=200 for all groups, Test statistic: *H*=89.58, *P*<0.001; Reject *H*_0_.

Steel-Dwass multiple comparisons: *P*<0.001 (Head vs. Body), *P*<0.001 (Head vs. Tail), *P*=0.71 (Body vs. Tail).

### ›Figure 5C

*H*_0_: The mean value of CITE in a certain time zone from the cell division is equal to zero.

*H_A_*: The mean value of CITE in a certain time zone from the cell division is not equal to zero.

*t* test, *n*=20 in the case of circumferential cell division and *n*=23 in that of longitudinal one,

Test statistic: absolute value of *t*: |*t*|

[Circ., 10-20 min]: |*t*|=0.97, *P*=0.34; Do not reject *H*_0_

[Circ., 20-30 min]: |*t*|=1.01, *P*=0.32; Do not reject *H*_0_

[Circ., 30-40 min]: |*t*|=1.12, *P*=0.29; Do not reject *H*_0_

[Circ., 40-50 min]: |*t*|=1.06, *P*=0.30; Do not reject *H*_0_

[Circ., 50-60 min]: |*t*|=2.37, *P*=0.03; Reject *H*_0_

[Circ., 60-70 min]: |*t*|=1.50, *P*=0.15; Do not reject *H*_0_

[Circ., 70-80 min]: |*t*|=1.09, *P*=0.29; Do not reject *H*_0_

[Circ., 80-90 min]: |*t*|=0.54, *P*=0.59; Do not reject *H*_0_

[Circ., 90-100 min]: |*t*|=1.75, *P*=0.10; Do not reject *H*_0_

[Circ., 100-110 min]: |*t*|=0.04, *P*=0.97; Do not reject *H*_0_

[Circ., 110-120 min]: |*t*|=0.35, *P*=0.73; Do not reject *H*_0_

[Long., 10-20 min]: |*t*|=2.02, *P*=0.06; Do not reject *H*_0_

[Long., 20-30 min]: |*t*|=0.51, *P*=0.61; Do not reject *H*_0_

[Long., 30-40 min]: |*t*|=1.05, *P*=0.30; Do not reject *H*_0_

[Long., 40-50 min]: |*t*|=0.98, *P*=0.34; Do not reject *H*_0_

[Long., 50-60 min]: |*t*|=0.21, *P*=0.83; Do not reject *H*_0_

[Long., 60-70 min]: |*t*|=1.03, *P*=0.31; Do not reject *H*_0_

[Long., 70-80 min]: |*t*|=1.08, *P*=0.29; Do not reject *H*_0_

[Long., 80-90 min]: |*t*|=0.31, *P*=0.76; Do not reject *H*_0_

[Long., 90-100 min]: |*t*|=0.03, *P*=0.97; Do not reject *H*_0_

[Long., 100-110 min]: |*t*|=1.75, *P*=0.09; Do not reject *H*_0_

[Long., 110-120 min]: |*t*|=0.24, *P*=0.81; Do not reject *H*_0_

### ›Figure 6D

*H*_0_: The mean values of tube diameter among the three groups of regime are all equal.

*H_A_*: The mean values of tube diameter among the three groups of regime are not all equal.

One-way analysis of variance, *n*=10 for all groups, *F*=791.10, *P*<0.001; Reject *H*_0_.

Tukey multiple comparisons: *P*<0.001 (No constriction vs. Mech-resp.), *P*<0.001 (No constriction vs. Scheduled), and *P*=0.002 (Mech-resp vs. Scheduled).

### ›Figure 6D´

*H*_0_: The mean values of curvedness among the three groups of regime are all equal.

*H_A_*: The mean values of curvedness among the three groups of regime are not all equal.

One-way analysis of variance, *n*=10 for all groups, *F*=575.25, *P*<0.001; Reject *H*_0_.

Tukey multiple comparisons: *P*<0.001 (No constriction vs. Mech-resp.), *P*<0.001 (No constriction vs. Scheduled), and *P*<0.001 (Mech-resp vs. Scheduled).

### ›Figure 7D´

*H*_0_: The aspect ratio of cells is the same in all three treatments of compression assay.

*H_A_*: The aspect ratio of cells is not the same in all three treatments of compression assay.

The Kruskal-Wallis test, *n*=500 for all groups, Test statistic: *H*=42.10, *P*<0.001; Reject *H*_0_.

Steelmultiplecomparisons:*P*<0.001(Controlvs.Lateral),*P*=0.043(Controlvs. Longitudinal).

### ›Figure 7D´´

*H*_0_: The angle of cells is the same in all three treatments of compression assay.

*H_A_*: The angle of cells is not the same in all three treatments of compression assay.

The Kruskal-Wallis test, *n*=500 for all groups, Test statistic: *H*=366.04, *P*<0.001; Reject *H*_0_.

Steelmultiplecomparisons:*P*<0.001(Controlvs.Lateral),*P*<0.001(Controlvs. Longitudinal).

### ›Figure 7E´

*H*_0_: The ratio of pMRLC to tMRLC is the same in all three treatments of compression assay.

*H_A_*: The ratio of pMRLC to tMRLC is not the same in all three treatments of compression assay.

The Friedman test, *n*=9 for all groups, Test statistic: *Q*=6, *P*=0.049; Reject *H*_0_.

Steelmultiplecomparisons:*P*=0.039(Controlvs.Lateral),*P*=1.82(Controlvs. Longitudinal).

### ›Figure 7E´

*H*_0_: The ratio of ROCK1 to tMRLC is the same in all three treatments of compression assay.

*H_A_*: The ratio of ROCK1 to tMRLC is not the same in all three treatments of compression assay.

The Friedman test, *n*=9 for all groups, Test statistic: *Q*=1.56, *P*=0.46; Do not reject *H*_0_.

Steel multiple comparisons: *P*=0.50 (Control vs. Lateral), *P*=0.60 (Control vs. Longitudinal).

**Figure S1.**
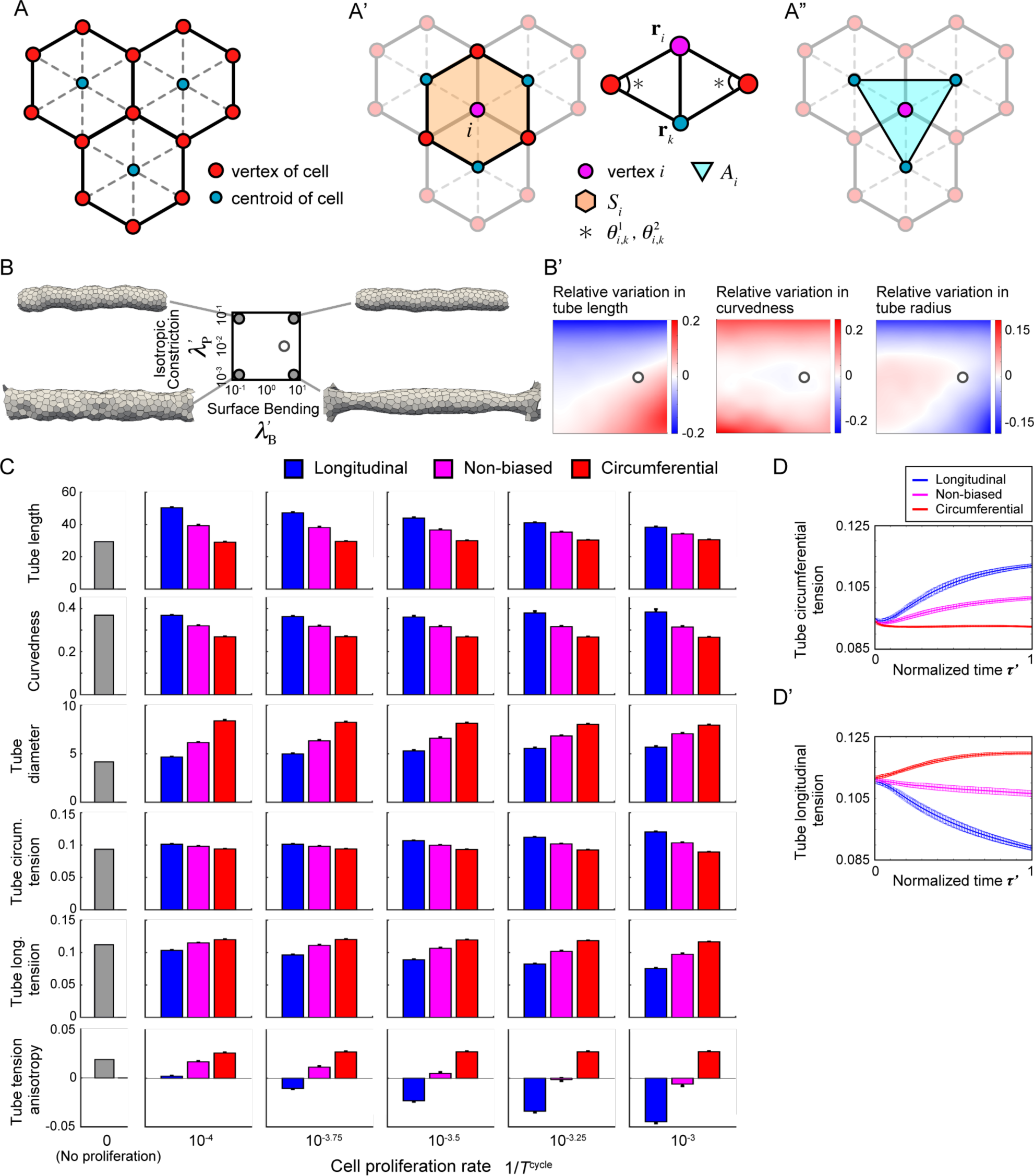
Preparation for the simulation without anisotropic junction constriction, corresponding to the Figure 2. (A-A”) Schematics for the explanation of discrete curvature in the vertex model. See the Materials and Methods (III-ii) for the notations. (B and B’) Diagram of generated tube morphology for parameters 
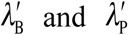
, and relative variation of morphological quantities in the absence of cell proliferation. Tubes are visualized in the four corners of parameter space (filled circles). Open circles represent parameter values used in this study. The axes labels are shown in B. (C) Parameter dependence of morphological and mechanical quantities for cell division orientation and cell proliferation rate. Color represents the type of cell division orientation. (D and D’) Time course of tube circumferential tension and that of tube longitudinal tension. See the Materials and Methods (III-vi) for the definition of tube circumferential/longitudinal tension.

**Figure S2.**
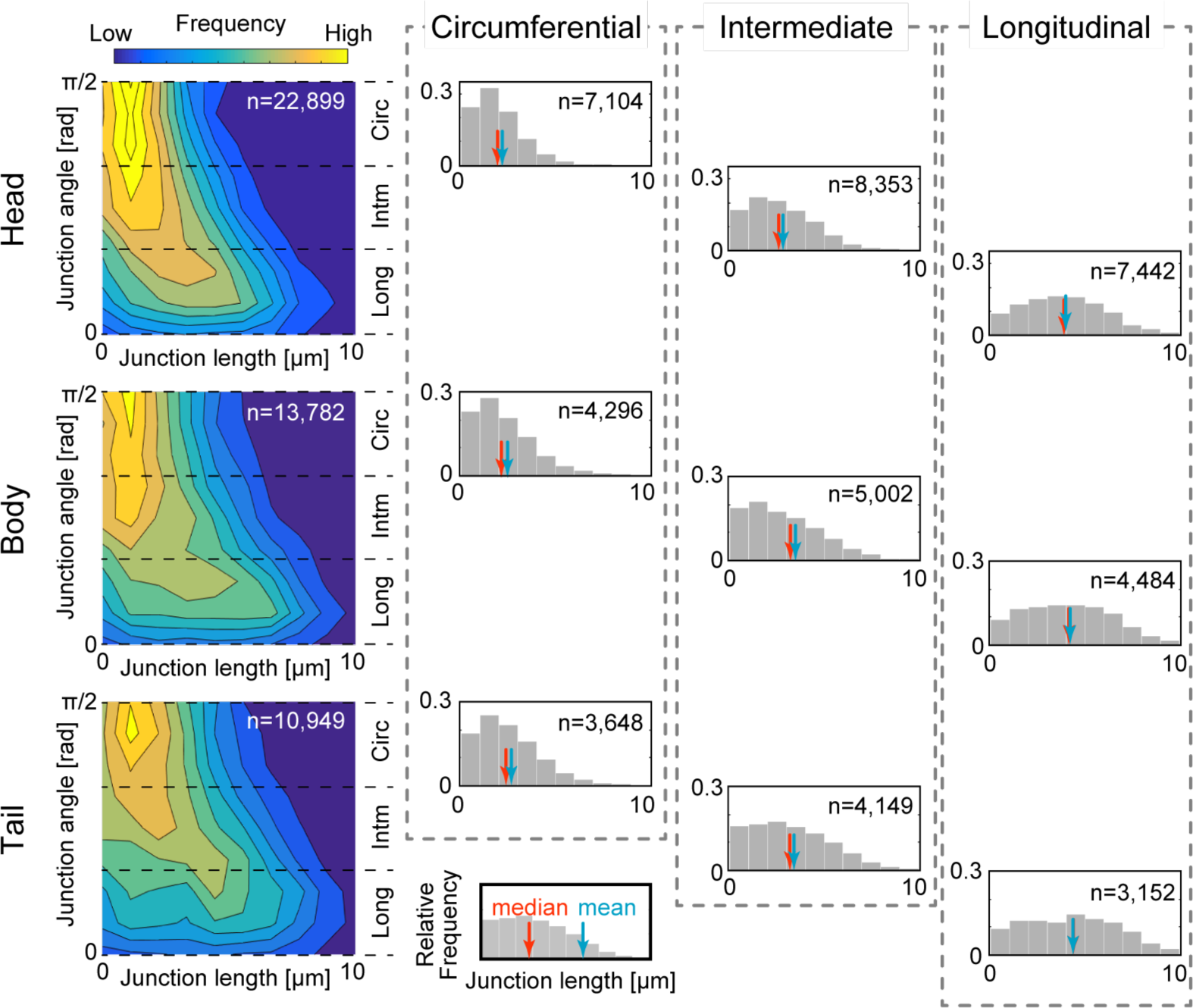
Apical junction length and angle, corresponding to the Figure 3. Frequency map on junction length *ℓ* and angle *θ* (left). Relative frequency in each region of epididymal tubule (Head, Body, and Tail) and angle category (Longitudinal, Intermediate, and Circumferential) is shown as histograms (right). The red/blue arrow indicates median/mean. The axis label of histograms is represented in the bottom. The circumferential junction length is shorter than the longitudinal junction length in any regions. In addition, the circumferential junction length in the head region is shorter than that in the tail region.

**Figure S3.**
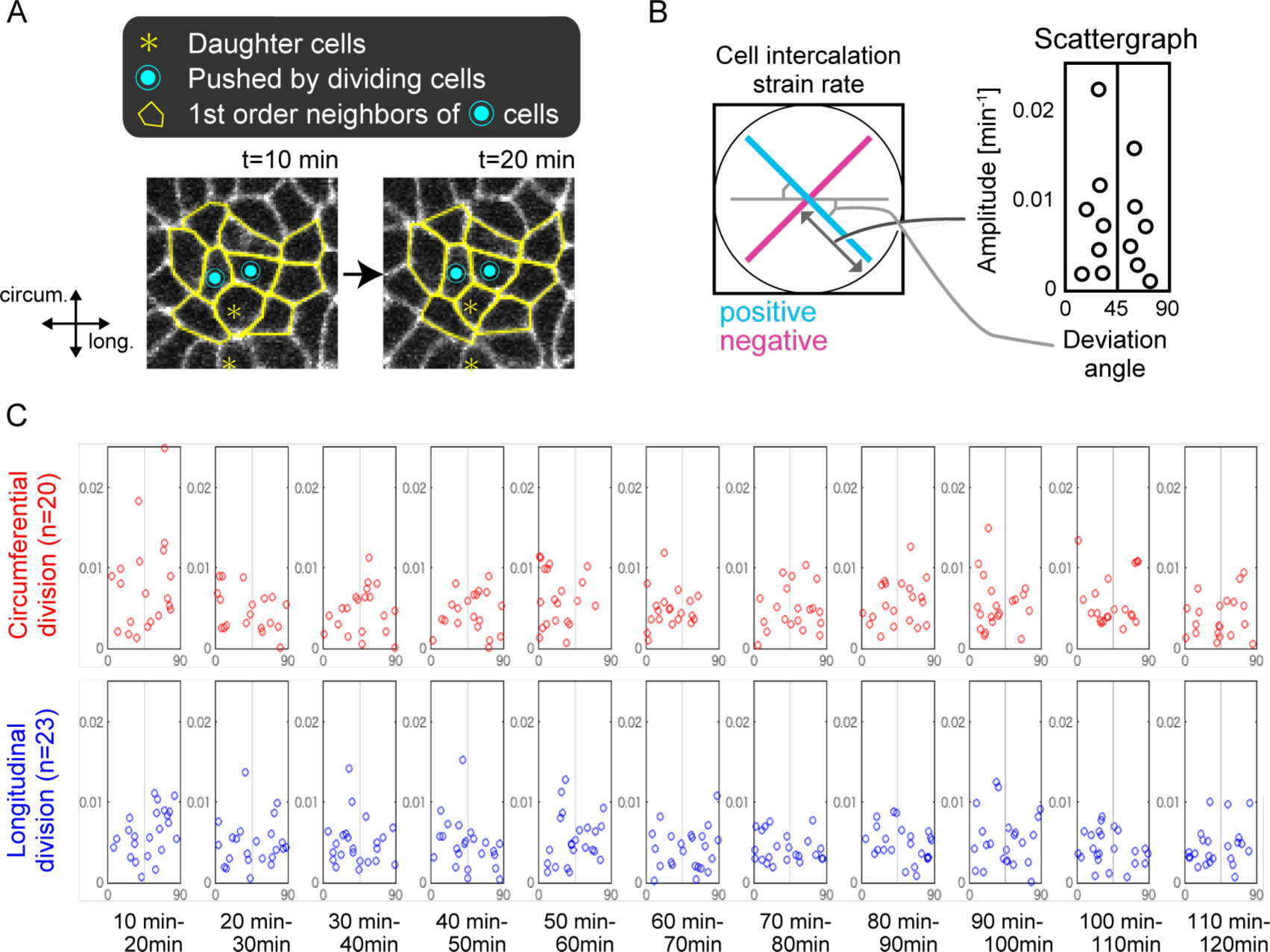
Measurement of cell intercalation strain rate from the live imaging data, corresponding to the Figure 5. (A) Illustration for target cells to calculate the cell intercalation strain rate. See the Materials and Methods (II-viii) for the details. (B) Graph representations for the cell intercalation strain rate. Orthogonal lines in the left graph represent principal strain rate. The amplitude and angle deviation of the positive principal strain rate (blue) in the left are expressed in the right angle-amplitude graph. (C) Time course of the cell intercalation strain rate as in the form of angle-amplitude graph for each type of cell division orientation: circumferential cell division and longitudinal one.

**Figure S4.**
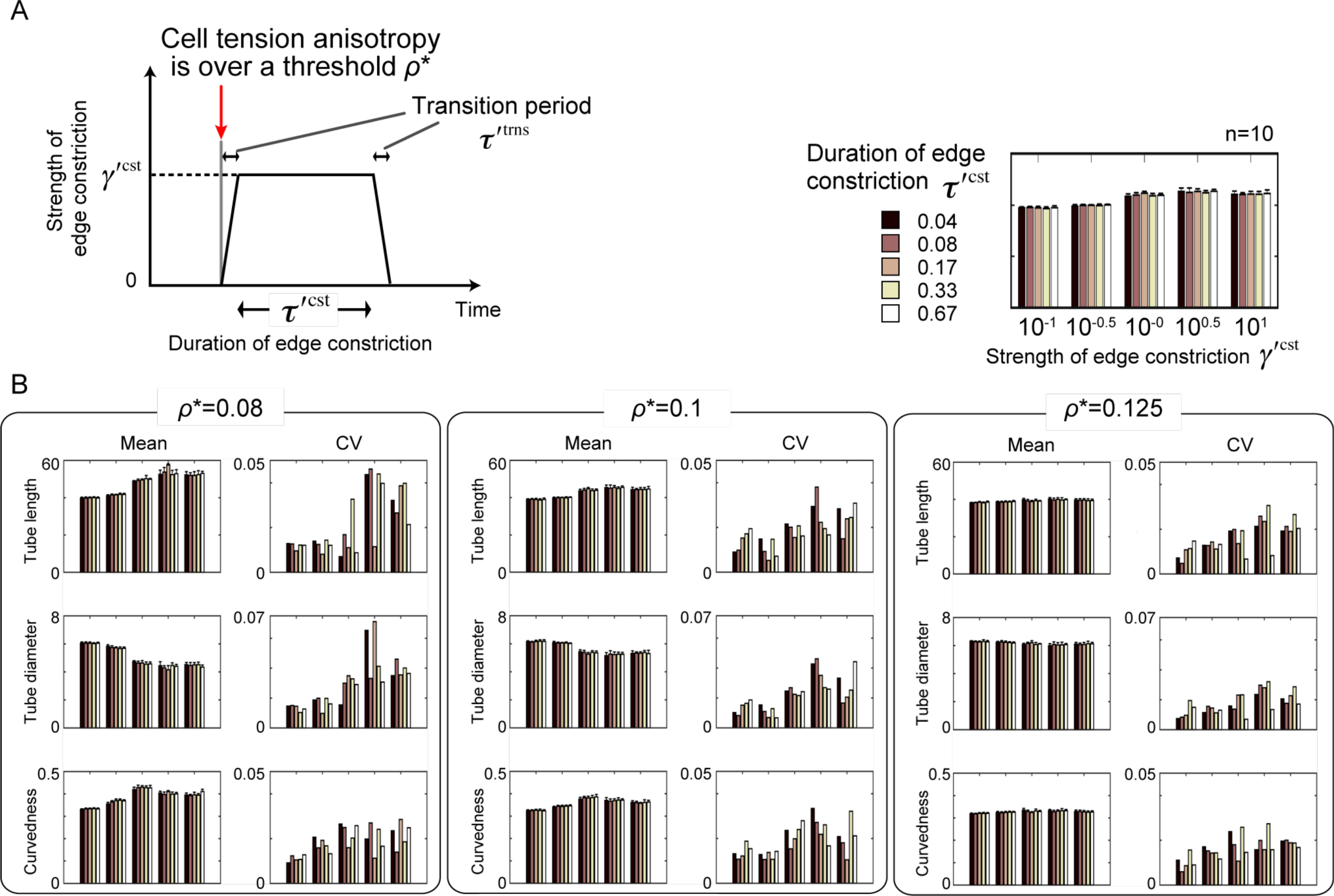
Preparation for the simulations in the mechano-responsive regime, corresponding to the Figure 6. (A) Schematics for the model. See the Materials and Methods (III-v) for the details. (B) Parameter dependence of tube morphology. The referenced graph is shown in the upper right. CV: coefficient of variance. See the Materials and Methods (III-vii) for determining the parameter values.

**Figure S5.**
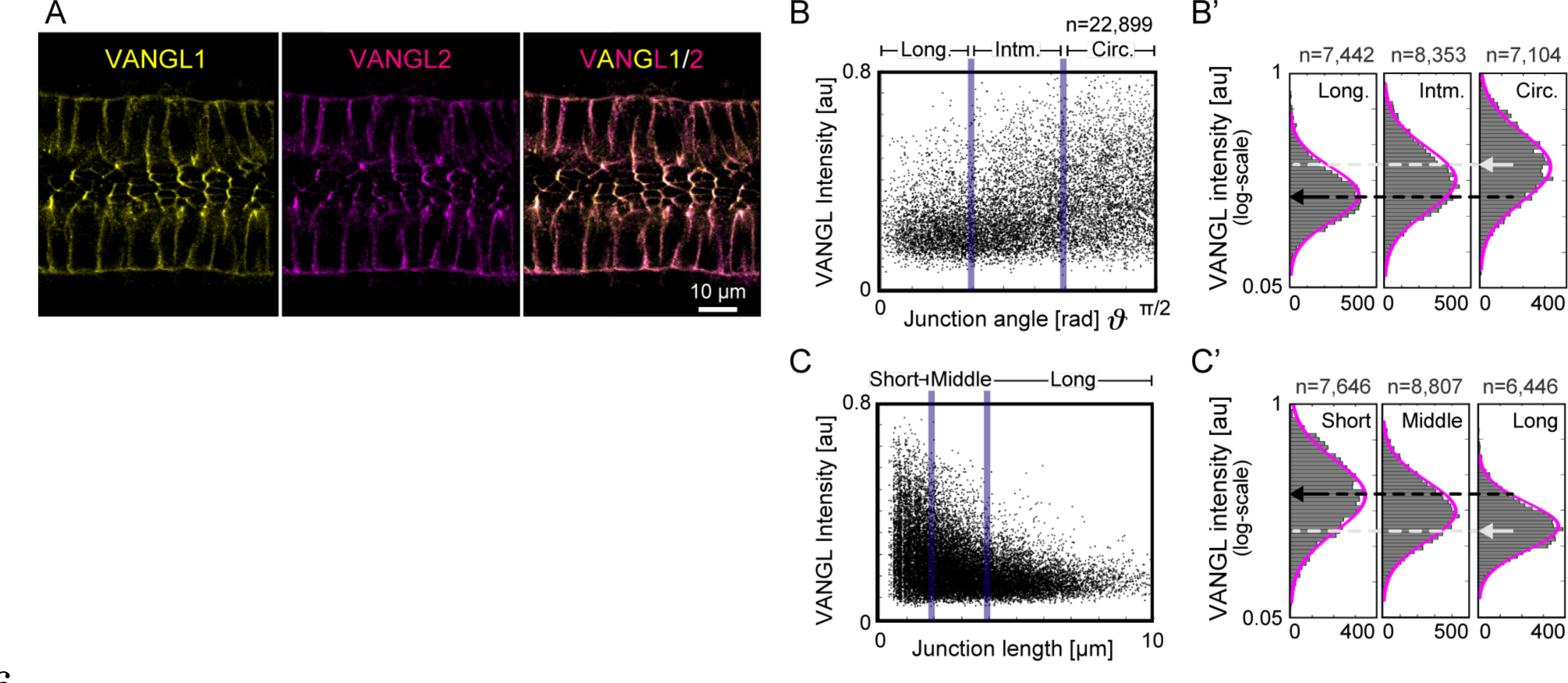
Figure S5 Spatial distribution of polarity proteins Van like protein (VANGL) 1 and 2, corresponding to the Discussion. (A) Immunofluorescence for VANGL1 and VANGL2 demonstrates that their co-localizations are on circumferential junctions of epididymal tubule. (B-C’) Relation of junction angle v.s. the fluorescence intensity, and that of junction length v.s. the intensity as with the form of Fig. 3E-F’. These graphs indicate the VANGL localization on the apical junctions of tubule shows significant polarization.

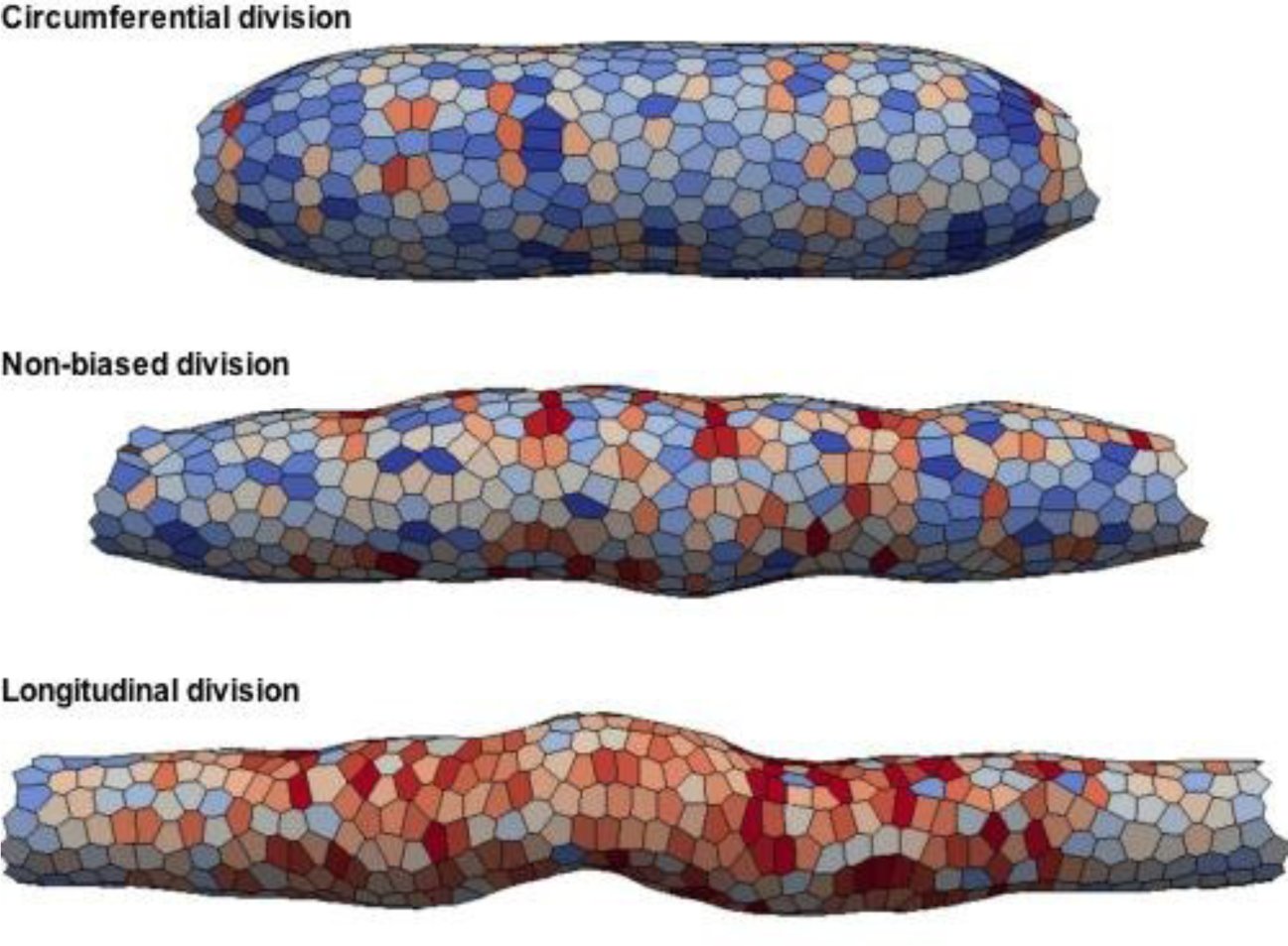
Movie 1 Model Simulation in Different Cell Division Orientation.

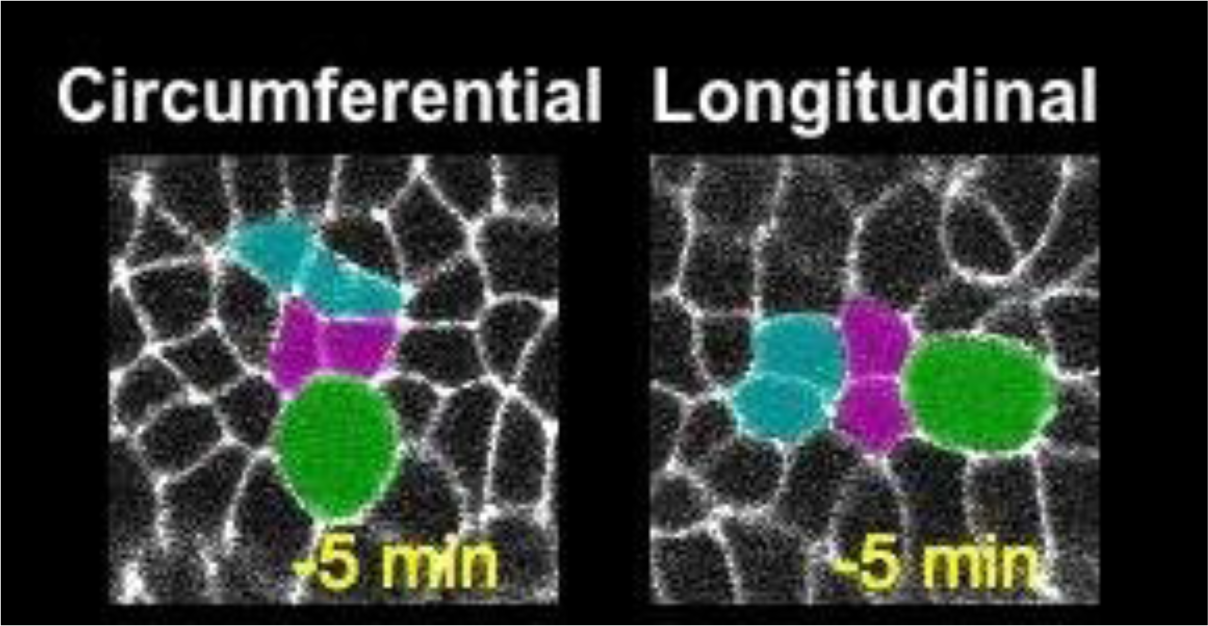
Movie 2 Live Imaging for Multicellular Behaviors against Cell Division.

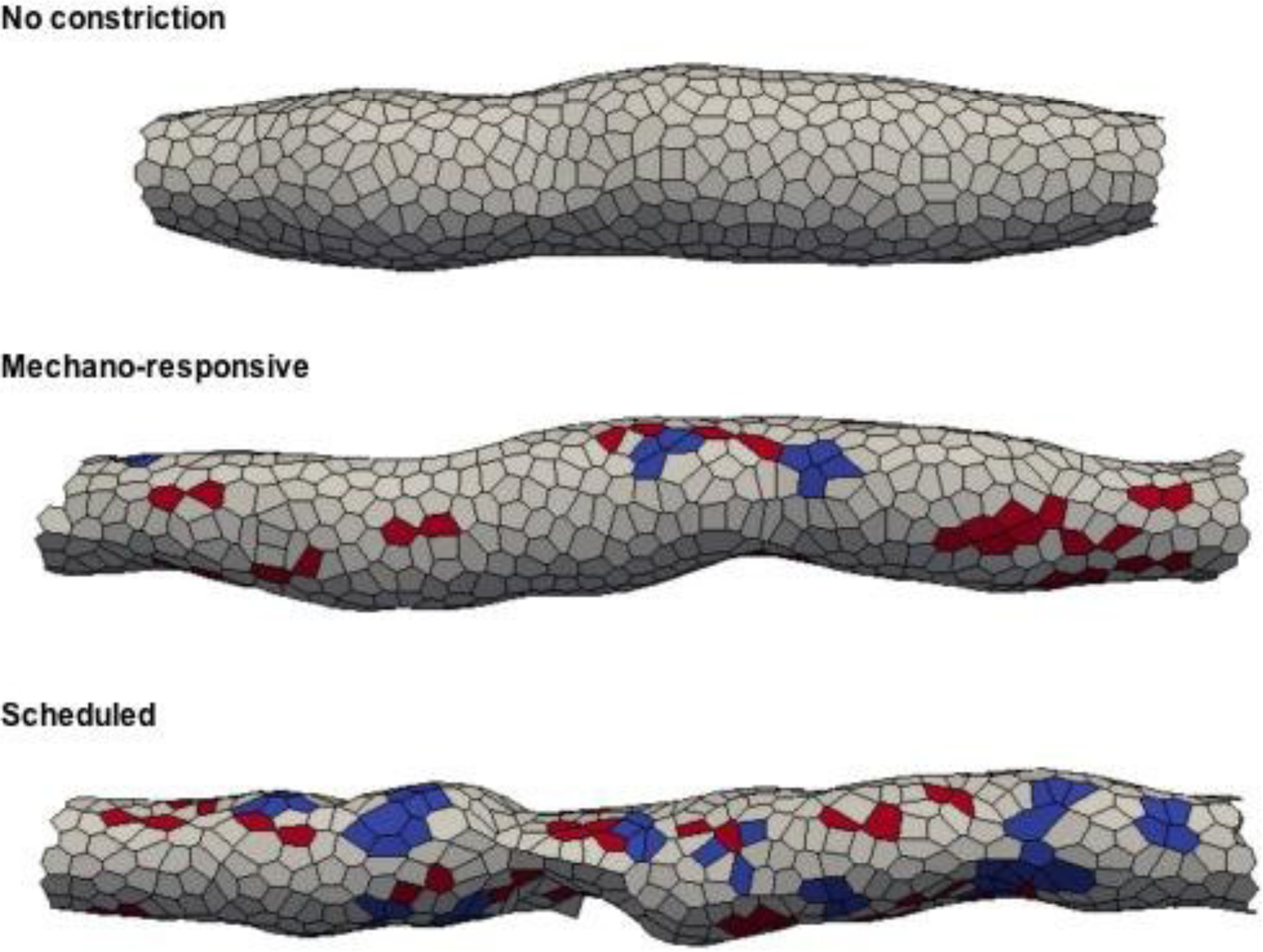
Movie 3 Model Simulation in Different Regimes.

## References

Abe, T., Kiyonari, H., Shioi, G., Inoue, K. I., Nakao, K., Aizawa, S. and Fujimori, T. (2011). Establishment of conditional reporter mouse lines at ROSA26 locus for live cell imaging. Genesis 49, 579–590.

Andrew, D. J. and Ewald, A. J. (2010). Morphogenesis of epithelial tubes: Insights into tube formation, elongation, and elaboration. Dev Biol 341, 34–55.

Batchelor, G. K. (1970). The stress system in a suspension of force-free particles. J. Fluid Mech 41, 545–570.

Beloussov, L. V (2015). Morphomechanics of Development. Springer.

Bertet, C., Sulak, L. and Lecuit, T. (2004). Myosin-dependent junction remodelling controls planar cell intercalation and axis elongation. Nature 429, 667–671.

Blanchard, G. B., Kabla, A. J., Schultz, N. L., Butler, L. C., Sanson, B., Gorfinkiel, N., Mahadevan, L. and Adams, R. J. (2009). Tissue tectonics: morphogenetic strain rates, cell shape change and intercalation. Nat. Methods 6, 458–464.

Bosveld, F., Bonnet, I., Guirao, B., Tlili, S., Wang, Z., Petitalot, A., Marchand, R., Bardet, P.-L., Marcq, P., Graner, F., et al. (2012). Mechanical Control of Morphogenesis by Fat/Dachsous/Four-Jointed Planar Cell Polarity Pathway. Science (80-.). 336, 724–727.

Collinet, C., Rauzi, M., Lenne, P. and Lecuit, T. (2015). Local and tissue-scale forces drive oriented junction growth during tissue extension. Nat. Cell Biol. 17, 1247–1258.

Dong, B., Hannezo, E. and Hayashi, S. (2014). Balance between Apical Membrane Growth and Luminal Matrix Resistance Determines Epithelial Tubule Shape. Cell Rep.

Du, X., Osterfield, M. and Shvartsman, S. Y. (2014). Computational analysis of three-dimensional epithelial morphogenesis using vertex models. Phys. Biol. 11, 66007.

Eisenhoffer, G. T., Loftus, P. D., Yoshigi, M., Otsuna, H., Chien, C.-B., Morcos, P. A. and Rosenblatt, J. (2012). Crowding induces live cell extrusion to maintain homeostatic cell numbers in epithelia. Nature 484, 546–549.

Farhadifar, R., Röper, J.-C., Aigouy, B., Eaton, S. and Jülicher, F. (2007). The influence of cell mechanics, cell-cell interactions, and proliferation on epithelial packing. Curr. Biol. 17, 2095–104.

Fischer, E., Legue, E., Doyen, A., Nato, F., Nicolas, J.-F., Torres, V., Yaniv, M. and Pontoglio, M. (2006). Defective planar cell polarity in polycystic kidney disease. Nat. Genet. 38, 21–23.

Fletcher, A. G., Osborne, J. M., Maini, P. K. and Gavaghan, D. J. (2013). Implementing vertex dynamics models of cell populations in biology within a consistent computational framework. Prog. Biophys. Mol. Biol. 113, 299–326.

Fletcher, A. G., Osterfield, M., Baker, R. E. and Shvartsman, S. Y. (2014). Vertex models of epithelial morphogenesis. Biophys. J. 106, 2291–2304.

Galkin, V. E., Orlova, A. and Egelman, E. H. (2012). Actin filaments as tension sensors. Curr. Biol. 22,.

Gudipaty, S. A., Lindblom, J., Loftus, P. D., Redd, M. J., Edes, K., Davey, C. F., Krishnegowda, V. and Rosenblatt, J. (2017). Mechanical stretch triggers rapid epithelial cell division through Piezo1. Nature 543, 118–121.

Guillot, C. and Lecuit, T. (2013). Mechanics of epithelial tissue homeostasis and morphogenesis. Science 340, 1185–9.

Hayakawa, K., Tatsumi, H. and Sokabe, M. (2011). Actin filaments function as a tension sensor by tension-dependent binding of cofilin to the filament. J. Cell Biol. 195, 721–727.

Hirashima, T. (2014). Pattern Formation of an Epithelial Tubule by Mechanical Instability during Epididymal Development. Cell Rep. 9, 866–73.

Hirashima, T. (2016). Mathematical study on robust tissue pattern formation in growing epididymal tubule. J. Theor. Biol. 407, 71–80.

Hirashima, T. and Adachi, T. (2015). Procedures for the Quantification of Whole-Tissue Immunofluorescence Images Obtained at Single-Cell Resolution during Murine Tubular Organ Development. PLoS One 10, e0135343.

Honda, H. and Eguchi, G. (1980). How much does the cell boundary contract in a monolayered cell sheet? J. Theor. Biol. 84, 575–588.

Honda, H., Tanemura, M. and Nagai, T. (2004). A three-dimensional vertex dynamics cell model of space-filling polyhedra simulating cell behavior in a cell aggregate. J. Theor. Biol. 226, 439–453.

Honda, H., Nagai, T. and Tanemura, M. (2008). Two different mechanisms of planar cell intercalation leading to tissue elongation. Dev. Dyn. 237, 1826–36.

Hufnagel, L., Teleman, A. A., Rouault, H., Cohen, S. M. and Shraiman, B. I. (2007). On the mechanism of wing size determination in fly development. Proc. Natl. Acad. Sci. U. S. A. 104, 3835–40.

Ishihara, S. and Sugimura, K. (2012). Bayesian inference of force dynamics during morphogenesis. J. Theor. Biol. 313, 201–211.

Joseph, A., Yao, H. and Hinton, B. T. (2009). Development and morphogenesis of the Wolffian/epididymal duct, more twists and turns. Dev Biol 325, 6–14.

Kantor, Y. and Nelson, D. R. (1987). Crumpling transition in polymerized membranes. Phys. Rev. Lett. 58, 2774–2777.

Kapur, J. N., Sahoo, P. K. and Wong, A. K. C. (1985). A new method for gray-level picture thresholding using the entropy of the histogram. Comput. Vision, Graph. Image Process. 29, 140.

Karner, C. M., Chirumamilla, R., Aoki, S., Igarashi, P., Wallingford, J. B. and Carroll, T. J. (2009). Wnt9b signaling regulates planar cell polarity and kidney tubule morphogenesis. Nat. Genet. 41, 793–799.

Kerschnitzki, M., Kollmannsberger, P., Burghammer, M., Duda, G. N., Weinkamer, R., Wagermaier, W. and Fratzl, P. (2013). Architecture of the osteocyte network correlates with bone material quality. J. Bone Miner. Res. 28, 1837–1845.

Kovács, M., Tóth, J., Hetényi, C., Málnási-Csizmadia, A. and Seller, J. R. (2004). Mechanism of blebbistatin inhibition of myosin II. J. Biol. Chem. 279, 35557–35563.

Kunimoto, K., Bayly, R. D., Vladar, E. K., Vonderfecht, T., Gallagher, A.-R. and Axelrod, J. D. (2017). Disruption of Core Planar Cell Polarity Signaling Regulates Renal Tubule Morphogenesis but Is Not Cystogenic. Curr. Biol.

Mammoto, T., Mammoto, A. and Ingber, D. E. (2013). Mechanobiology and Developmental Control. Annu. Rev. Cell Dev. Biol. 29, 27–61.

Mao, Y., Tournier, A. L., Hoppe, A., Kester, L., Thompson, B. J. and Tapon, N. (2013). Differential proliferation rates generate patterns of mechanical tension that orient tissue growth. EMBO J. 32, 2790–2803.

Meyer, M., Desbrun, M., Schr, P. and Barr, A. H. (2003). Discrete Differential-Geometry Operators for Triangulated 2-Manifolds. Vis. Math. III 113–134.

Morin, X. and Bellaïche, Y. (2011). Mitotic Spindle Orientation in Asymmetric and Symmetric Cell Divisions during Animal Development. Dev. Cell 21, 102–119.

Nagai, T. and Honda, H. (2001). A dynamic cell model for the formation of epithelial tissues. Philos. Mag. Part B 81, 699–719.

Nishimura, T., Honda, H. and Takeichi, M. (2012). Planar cell polarity links axes of spatial dynamics in neural-tube closure. Cell 149, 1084–1097.

Ohyama, T. and Groves, A. K. (2004). Generation of Pax2-Cre Mice by Modification of a Pax2 Bacterial Artificial Chromosome. Genesis 38, 195–199.

Okuda, S., Inoue, Y., Eiraku, M., Sasai, Y. and Adachi, T. (2013). Reversible network reconnection model for simulating large deformation in dynamic tissue morphogenesis. Biomech. Model. Mechanobiol. 12, 627–644.

Okuda, S., Inoue, Y., Eiraku, M., Adachi, T. and Sasai, Y. (2014). Vertex dynamics simulations of viscosity-dependent deformation during tissue morphogenesis. Biomech. Model. Mechanobiol. 14, 413–425.

Osterfield, M., Du, X., Schüpbach, T., Wieschaus, E. and Shvartsman, S. Y. (2013). Three-Dimensional Epithelial Morphogenesis in the Developing Drosophila Egg. Dev. Cell 24, 400–410.

Rauzi, M., Verant, P., Lecuit, T. and Lenne, P.-F. (2008). Nature and anisotropy of cortical forces orienting Drosophila tissue morphogenesis. Nat. Cell Biol. 10, 1401–10.

Sasai, Y. (2013). Cytosystems dynamics in self-organization of tissue architecture. Nature 493, 318–26.

Savin, T., Kurpios, N. A., Shyer, A. E., Florescu, P., Liang, H., Mahadevan, L. and Tabin, C. J. (2011). On the growth and form of the gut. Nature 476, 57–62.

Seung, H. S. and Nelson, D. R. (1988). Defects in flexible membranes with crystalline order. Phys. Rev. A 38, 1005–1018.

Shraiman, B. I. (2005). Mechanical feedback as a possible regulator of tissue growth. Proc. Natl. Acad. Sci. U. S. A. 102, 3318–3323.

Stewart, M. P., Helenius, J., Toyoda, Y., Ramanathan, S. P., Muller, D. J. and Hyman, A. a (2011). Hydrostatic pressure and the actomyosin cortex drive mitotic cell rounding. Nature 469, 226–230.

Sugimura, K. and Ishihara, S. (2013). The mechanical anisotropy in a tissue promotes ordering in hexagonal cell packing. Development 140, 4091–4101.

Susaki, E. A., Tainaka, K., Perrin, D., Kishino, F., Tawara, T., Watanabe, T. M., Yokoyama, C., Onoe, H., Eguchi, M., Yamaguchi, S., et al. (2014). Whole-brain imaging with single-cell resolution using chemical cocktails and computational analysis. Cell 157, 726–739.

Tang, N., Marshall, W. F., McMahon, M., Metzger, R. J. and Martin, G. R. (2011). Control of Mitotic Spindle Angle by the RAS-Regulated ERK1/2 Pathway Determines Lung Tube Shape. Science (80-.). 333, 342–345.

Thompson, D. W. (1942). On growth and form.

Tomaszewski, J., Joseph, A., Archambeault, D. and Yao, H. H. (2007). Essential roles of inhibin beta A in mouse epididymal coiling. Proc Natl Acad Sci U S A 104, 11322–11327.

Walck-Shannon, E. and Hardin, J. (2014). Cell intercalation from top to bottom. Nat. Rev. Mol. Cell Biol. 15, 34–48.

Watanabe, T., Hosoya, H. and Yonemura, S. (2007). Regulation of Myosin II Dynamics by Phosphorylation and Dephosphorylation of Its Light Chain in Epithelial Cells. Mol. Biol. Cell 18, 605–616.

Xu, B., Washington, A. M., Domeniconi, R. F., Ferreira Souza, A. C., Lu, X., Sutherland, A. and Hinton, B. T. (2016). Protein tyrosine kinase 7 is essential for tubular morphogenesis of the Wolffian duct. Dev. Biol. 412, 219–233.

Yokomizo, T., Yamada-Inagawa, T., Yzaguirre, A. D., Chen, M. J., Speck, N. A. and Dzierzak, E. (2012). Whole-mount three-dimensional imaging of internally localized immunostained cells within mouse embryos. Nat. Protoc. 7, 421–431.

Zallen, J. A. and Blankenship, J. T. (2008). Multicellular dynamics during epithelial elongation. Semin. Cell Dev. Biol. 19, 263–270.

Zar, J. H. (2014). Biostatistical Analysis, 5th Edition.

